# A combined DTI-fMRI approach for optimizing the delineation of posteromedial vs. anterolateral entorhinal cortex

**DOI:** 10.1101/2022.12.23.520976

**Authors:** Ingrid Framås Syversen, Daniel Reznik, Menno P. Witter, Asgeir Kobro-Flatmoen, Tobias Navarro Schröder, Christian F. Doeller

**Author notes:** Corresponding author. Postal address: Kavli Institute for Systems Neuroscience, MH, NTNU, Postbox 8905, 7491 Trondheim, Norway. Present address: Akershus University Hospital, Postbox 1000, 1478 Nordbyhagen, Norway (Ingrid Framås Syversen).

## Abstract

In the entorhinal cortex (EC), attempts have been made to identify the human homologue regions of the medial (MEC) and lateral (LEC) subdivision using either functional magnetic resonance imaging (fMRI) or diffusion tensor imaging (DTI). However, there are still discrepancies between entorhinal subdivisions depending on the choice of connectivity seed regions and the imaging modality used. While DTI can be used to follow the white matter tracts of the brain, fMRI can identify functionally connected brain regions. In this study, we used both DTI and resting-state fMRI in 103 healthy adults to investigate both structural and functional connectivity between the EC and associated cortical brain regions. Differential connectivity with these regions was then used to predict the locations of the human homologues of MEC and LEC. Our results from combining DTI and fMRI support a subdivision into posteromedial (pmEC) and anterolateral (alEC) EC and reveal a confined border between the pmEC and alEC. Furthermore, the EC subregions obtained by either imaging modality showed similar distinct connectivity patterns: While pmEC showed increased connectivity preferentially with the default mode network, the alEC exhibited increased connectivity with regions in the dorsal attention and salience networks. Optimizing the delineation of the human homologues of MEC and LEC with a combined, cross-validated DTI-fMRI approach allows to define a likely border between the two subdivisions and has implications for both cognitive and translational neuroscience research.

## 1 Introduction

The entorhinal cortex (EC) is part of the medial temporal lobe (MTL), the key episodic memory system of the mammalian brain (Amaral et al., 1987). In rodents, two main subregions of the EC have been defined cytoarchitectonically – the ‘medial’ entorhinal cortex (MEC) and ‘lateral’ entorhinal cortex (LEC) – which differ both in their functional properties and connectivity to other brain regions (Canto et al., 2008; Kerr et al., 2007; Knierim et al., 2014; Nilssen et al., 2019; Ranganath and Ritchey, 2012; van Strien et al., 2009). The EC is central for memory formation, spatial navigation and time perception, and MEC and LEC are traditionally viewed as being part of separate information streams related to these different processes (Eichenbaum et al., 2007; Moser and Moser, 2013; Tsao et al., 2018). Precise delineations of the human homologues of the rodent MEC and LEC are missing, though studies using either functional magnetic resonance imaging (fMRI) or diffusion tensor imaging (DTI) support a subdivision into an entorhinal posteromedial (pmEC) vs. anterolateral (alEC) part (Maass et al., 2015; Navarro Schröder et al., 2015; Reagh and Yassa, 2014; Schultz et al., 2012; Syversen et al., 2021). However, these studies show different extents of subdivisions along the posterior-anterior (PA) vs. medial-lateral (ML) axes, and it is still unclear if and how estimates of connectional properties of the subregions correspond or differ across MRI modalities (structural DTI and functional MR imaging, respectively) within the same individuals.

Traditionally, the EC has been viewed as the main interface between the neocortex and the hippocampus (Buzsáki, 1996; Lavenex and Amaral, 2000), processing and relaying information in two separate information streams (Diana et al., 2007; Eichenbaum et al., 2007; Nilssen et al., 2019; Ranganath and Ritchey, 2012; Witter et al., 2017). In this dual stream model, the MEC is involved in allocentric processing of space (the “where” pathway), while the LEC is involved in processing of objects and time (the “what” pathway). Two task-based human fMRI studies investigating spatial (scenes) vs. non-spatial (objects) processing found differences between medial and lateral parts of the EC, respectively (Reagh and Yassa, 2014; Schultz et al., 2012). Another study, however, reported similar functional differences between posteromedial and anterolateral EC (Navarro Schröder et al., 2015). Furthermore, the dual stream model implies that there are separate connectivity pathways between parahippocampal cortex (PHC) and the human homologue of MEC, and between perirhinal cortex (PRC) and the human homologue of LEC. One of the fMRI studies subdividing the EC into pmEC and alEC (Maass et al., 2015) was based on this assumption. However, new evidence obtained in rodents shows that the postrhinal cortex (which corresponds to PHC in humans) in fact projects at least as much to MEC as to LEC (Doan et al., 2019; Nilssen et al., 2019). Recent DTI and fMRI results from humans furthermore suggest extensive direct connections between the neocortex and the hippocampus and subicular complex bypassing the EC and that the connections are less hierarchical and segregated than previously assumed (Dalton et al., 2022; Grande et al., 2022; Huang et al., 2021; Ma et al., 2022; Reznik et al., 2023; Rolls et al., 2022). Results that were stringently based on the old dual stream model should therefore be re-evaluated in light of these new findings.

It is furthermore not clear whether the differences in previous segmentation results originate solely from using different seed regions, or if differences in imaging modalities might also affect proposed divisional schemes for EC. While fMRI investigates functional connectivity (Smitha et al., 2017; Van Dijk et al., 2010), DTI identifies structural connectivity paths (Jeurissen et al., 2019; Mori and Zhang, 2006). Functionally connected brain regions are involved in the same processing, and although function generally is constrained by anatomy, it does not necessarily mean that functionally connected regions are monosynaptically connected (Rykhlevskaia et al., 2008). Conversely, brain regions do not always have to be involved in the same functional processes even though they are structurally connected. It can therefore be difficult to quantitatively compare structural and functional connectivity. Previous fMRI studies subdivided the EC into pmEC and alEC based on functional connectivity with the PHC vs. PRC as well as with distal vs. proximal subiculum, and with posterior-medial vs. anterior-temporal cortical systems (shown to be functionally distinct, see Ranganath and Ritchey (2012)), respectively (Maass et al., 2015; Navarro Schröder et al., 2015). Our recent DTI study aimed to identify the human homologue regions of MEC and LEC using differential EC structural connectivity, and thus segregated EC connectivity with the presubiculum and retrosplenial cortex (RSC) on the one hand, and with the distal CA1, the proximal subiculum (dCA1pSub), and the posterolateral orbitofrontal cortex (OFC) on the other hand (Syversen et al., 2021). These seed regions were chosen because of their established connections with MEC and LEC in rodents (Caballero-Bleda and Witter, 1993; Honda and Ishizuka, 2004; Hoover and Vertes, 2007; Insausti and Amaral, 2008; Jones and Witter, 2007; Kondo and Witter, 2014; Saleem et al., 2008; Syversen et al., 2021; Witter and Amaral, 1991; Witter and Amaral, 2021; Wyss and Van Groen, 1992), and thus to segment the EC based on connectivity both with hippocampal (presubiculum, dCA1pSub) and neocortical (RSC, OFC) regions. This resulted in a subdivision of the EC into putative pmEC vs. alEC, although with some differences in medial-lateral (ML) vs. posterior-anterior (PA) orientation of the border between the subregions compared to the abovementioned fMRI segmentations (Syversen et al., 2021).

In the present study, we used both DTI and fMRI data acquired in the same cohort of participants to perform structural and functional connectivity analysis between the EC and selected brain regions. First, we aimed to replicate our previous DTI results (Syversen et al., 2021). Then, we compared structural connectivity from DTI with functional connectivity from resting-state (rs-)fMRI between the EC and selected seed regions. Ultimately, this allowed us to use a combination of structural and functional connectivity results to predict the locations of the human homologues of MEC and LEC. Following new insights from rodents, MEC is here defined to be differentially connected with presubiculum and RSC, whereas LEC is defined to be differentially connected with dCA1pSub and OFC (Caballero-Bleda and Witter, 1993; Honda and Ishizuka, 2004; Hoover and Vertes, 2007; Insausti and Amaral, 2008; Jones and Witter, 2007; Kondo and Witter, 2014; Saleem et al., 2008; Syversen et al., 2021; Witter and Amaral, 1991; Witter and Amaral, 2021; Wyss and Van Groen, 1992). The overarching goal of the study was to extend and bridge the results from previous studies where DTI and fMRI has been investigated separately (Maass et al., 2015; Navarro Schröder et al., 2015; Syversen et al., 2021), in order to obtain a high-certainty, multimodal definition of the human homologues of rodent MEC and LEC.

## 2 Materials and methods

### 2.1 MRI data

Structural, diffusion-weighted and resting-state functional MRI data from 184 healthy adults were obtained from the WU-Minn Human Connectome Project (HCP; http://db.humanconnectome.org), in line with the WU-Minn HCP Consortium Open Access Data Use Terms (Marcus et al., 2011; Van Essen et al., 2012). All participants provided written informed consent, and the study was approved by the Institutional Review Board of Washington University in St. Louis, MO, USA. Detailed image acquisition protocols are provided in the HCP Reference Manual (https://humanconnectome.org/storage/app/media/documentation/s1200/HCP_S1200_Relea <u>se_Reference_Manual.pdf</u>). In short, 3T MRI data were acquired on a Siemens Connectome Skyra scanner, while 7T MRI data were acquired on a Siemens Magnetom scanner. Structural T1-weighted images were acquired at 3T using a 3D MPRAGE sequence with 0.7 mm isotropic resolution. Diffusion-weighted images were acquired at 3T and 7T, using spin-echo EPI sequences with 1.25 and 1.05 mm isotropic resolution, respectively, and with b-values of 1000, 2000, 3000 s/mm^2^ and 1000, 2000 s/mm^2^ in addition to a set of b = 0 images (Feinberg et al., 2010; Sotiropoulos et al., 2013). fMRI data were acquired at 7T using a gradient-echo EPI sequence with 1.6 mm isotropic resolution and a TR of 1000 ms (Moeller et al., 2010; Setsompop et al., 2012; Xu et al., 2012). There were two resting-state runs with posterior-anterior (PA) and two runs with anterior-posterior (AP) phase encoding direction, and in each run 900 image volumes were acquired over 16 minutes.

### 2.2 Preprocessing

The data were minimally preprocessed by the HCP processing pipeline as described in detail in Glasser et al. (2013). In brief, the processing included defacing of structural MR images, automated cortical parcellation and brain extraction, and calculation of the registration transformations between the participants’ native structural space and MNI space (Fischl, 2012; Jenkinson et al., 2002; Jenkinson et al., 2012; Milchenko and Marcus, 2013). DTI and fMRI data were corrected for gradient nonlinearity, motion and geometric distortions, in addition to Eddy current correction for DTI and denoising for fMRI (Andersson et al., 2003; Andersson and Sotiropoulos, 2015; Andersson and Sotiropoulos, 2016). The preprocessed and denoised fMRI data were provided in MNI space by the HCP, while DTI data remained in the participants’ native space (although rigidly aligned to the structural images).

Because the HCP data were not specifically optimized for image quality in the medial temporal lobe, careful quality control was performed. Eighty-one participants were excluded due to insufficient quality control measures, including motion estimates (n = 6), temporal signal-to-noise ratio (tSNR; n = 30) and missing data (e.g. missing structural data, transforms, preprocessing etc.; n = 45). Exclusion criteria for fMRI data were maximum absolute root-mean-square motion ≥ 2 mm during the whole run, and slice tSNR ≤ 150 (see Supplementary Figure 1). The remaining 103 participants with both DTI and rs-fMRI data were included for further analyses (note that included participants could have some excluded rs-fMRI runs, but had complete DTI data and at least one included rs-fMRI run). In total, 3T and 7T DTI data for all 103 participants were included, in addition to 171 rs-fMRI runs with PA phase encoding direction and 139 rs-fMRI runs with AP phase encoding direction.

#### 2.2.1 Registration

To facilitate group analysis and comparison with results from our previous DTI study (Syversen et al., 2021), the registration transform between the participants’ HCP MNI space and the MNI152-09b template (Fonov et al., 2009) was determined, using symmetric non-linear registration in the Advanced Neuroimaging Toolbox (ANTs; version 2.3.4, http://stnava.github.io/ANTs/) based on mutual information (Avants et al., 2011). DTI data in the participants’ native space could thus be registered to the version of MNI space used by the HCP using the transform they provided, and fMRI and DTI data could then be further registered to MNI152-09b space using the transform obtained from ANTs.

#### 2.2.2 Regions of interest

Regions of interest (ROIs) were extracted from the automated cortical parcellations obtained from the FreeSurfer (version 7.1.1, https://surfer.nmr.mgh.harvard.edu/) functions *recon-all* and *segmentHA_T1* on the MNI152-09b template (Fischl et al., 2002; Fischl et al., 2004; Iglesias et al., 2015). In particular, template ROIs of the EC, presubiculum, distalCA1 + proximal subiculum (dCA1pSub), RSC and OFC were defined as in our previous work (Syversen et al., 2021): The EC ROI from FreeSurfer was considered to extend too far posteriorly towards the parahippocampal cortex and laterally beyond the collateral sulcus, and was therefore masked by a probabilistic EC ROI from the Jülich-Brain Cytoarchitectonic Atlas thresholded at 0.25 (Amunts et al., 2020). The resulting EC ROI was further refined by using the FMRIB Software Library’s (FSL; version 5.0.9, http://fsl.fmrib.ox.ac.uk/fsl/) function *fslmaths-ero* to erode the ROI once, before manually removing remaining voxels creating discontinuities of the surface of the ROI in posterior and lateral parts (Supplementary Figure 2A). The presubiculum ROI from FreeSurfer was used as is. ROIs of distal CA1 and proximal subiculum were created by splitting the ROIs of each of the two hippocampal structures in half along their proximodistal axis. That is, of all voxels encompassing CA1, the half located distally was included, and of all the voxels encompassing subiculum, the half located proximally was included – these two halves thus make up what we here define and refer to as ‘distal CA1 + proximal subiculum’ (dCA1pSub). The body and the head of the dCA1+pSub were used, but the parts with folding were removed from the head because of its complicated geometry. To define RSC and OFC ROIs, respectively, the FreeSurfer parcellations named “isthmus cingulate” and “lateral orbitofrontal” were used as starting points. The final RSC ROI was obtained by tailoring the isthmus cingulate and removing the excess superior areas (Supplementary Figure 2B), whereas the final OFC ROI was obtained by extracting the posterolateral quadrant of the lateral orbitofrontal area. All resulting template ROIs can be seen in Supplementary Figure 3. These MNI152-09b template ROIs were then registered to the participants’ individual spaces and further masked by the corresponding individual parcellations, in order to increase individual anatomical precision of the ROI boundaries.

### 2.3 DTI analysis

Voxel-wise fiber orientation distribution functions (fODFs) for the 3T DTI data were provided by the HCP. For the 7T DTI data, fODFs were computed in the same manner using FSL’s *bedpostx* (Hernández et al., 2013; Sotiropoulos et al., 2016). Probabilistic tractography between ROIs was performed by running FSL’s *probtrackx2* separately on the fODFs from 3T and 7T (Behrens et al., 2007; Behrens et al., 2003b; Hernandez-Fernandez et al., 2019). Tractography was run in ROI-by-ROI mode to visualize the structural connectivity paths between the ROIs, and in voxel-by-ROI mode to create connectivity maps from the EC ROI to the other ROIs. For more details about parameters used for *bedpostx* and *probtrackx2*, see Syversen et al. (2021). All tractography results were registered to MNI152-09b space and all further analyses were performed there to facilitate group analyses.

### 2.4 fMRI analysis

Seed-based functional connectivity analysis was performed on the rs-fMRI data using an in-house MATLAB script (version R2020b, MathWorks, Natick, MA, USA). All fMRI volumes were smoothed with a 6 mm full-width half-maximum (FWHM) Gaussian kernel. This kernel was chosen empirically after testing several different kernel widths (3-6 mm), where using smoothing kernels smaller than 6 mm generally showed weaker whole-brain functional connectivity results. For each included rs-fMRI run and each ROI, Pearson correlation was calculated between the time series of the average ROI signal and all the other voxels in the brain to create whole-brain functional connectivity maps. The resulting maps were Fisher-Z transformed and registered to MNI152-09b space to facilitate group analyses and comparison.

### 2.5 Group analysis

Group averages of EC structural connectivity maps (from DTI analysis) and whole-brain functional connectivity maps (from fMRI analysis) were created by adding together the results from all included participants, both separately and across field strengths for DTI and phase encoding directions for rs-fMRI. For rs-fMRI, EC maps of functional connectivity with the other ROIs were created by masking the group-averaged whole-brain functional connectivity maps by an EC ROI.

#### 2.5.1 MEC and LEC segmentation

Group-averaged EC structural and functional connectivity maps were used to segment the EC into respective DTI-based and rs-fMRI-based MEC and LEC homologues. MEC was defined as being preferentially connected with presubiculum and/or RSC, whereas LEC was defined as being preferentially connected with dCA1pSub and/or OFC (Caballero-Bleda and Witter, 1993; Honda and Ishizuka, 2004; Hoover and Vertes, 2007; Insausti and Amaral, 2008; Jones and Witter, 2007; Kondo and Witter, 2014; Saleem et al., 2008; Syversen et al., 2021; Witter and Amaral, 1991; Witter and Amaral, 2021; Wyss and Van Groen, 1992). These regions were chosen because both MEC and LEC would then be defined by one neocortical region (RSC and OFC, respectively) and one region within the hippocampal formation (presubiculum and dCA1pSub, respectively). The segmentation of the EC was performed using FSL’s *find_the_biggest* (Behrens et al., 2003a; Johansen-Berg et al., 2004) on the connectivity maps. Intermediate analyses showed, however, that the connectivity values varied substantially between maps from different ROIs, which produced MEC/LEC segmentations of very unequal sizes (see Supplementary Table 1). To ensure more balanced sizes of resulting MECs and LECs, the connectivity maps were therefore iteratively scaled (up or down; constrained by the boundaries of the EC ROI) until the MEC/LEC size ratio was in the range [0.95,1.05] for each hemisphere separately (Supplementary Figure 4). Segmentation was performed separately for field strengths for DTI and phase encoding directions for rs-fMRI, based on the 2×2 different combinations of seed ROIs in addition to a combination of ROIs (presubiculum+RSC and dCA1pSub+OFC), resulting in a total of 20 different segmentations (see Supplementary Table 2 for an overview). To create final MEC and LEC segmentations for DTI and rs-fMRI, the connectivity maps from the combination of ROIs were averaged across field strengths for DTI and phase encoding directions for rs-fMRI, before performing the same segmentation process as described above.

#### 2.5.2 Segmentation comparisons

To determine whether the predicted MEC and LEC homologues were located primarily along a posterior-anterior (PA) or a medial-lateral (ML) axis, the degree of PA and ML orientation of the border between the subregions was calculated for all segmentation approaches (Syversen et al., 2021). This was defined as a percentage between 0 and 100%, where a 100% PA orientation of the border would mean that the MEC and LEC homologues were located strictly posteriorly-anteriorly with respect to each other, and a 100% ML orientation would mean that they were located strictly medially-laterally with respect to each other. The percentage was calculated by determining the angle between the vector from the MEC to LEC centers of gravity, and a vector corresponding to a pure PA or ML axis. Furthermore, to determine the correspondence between the final DTI and rs-fMRI segmentations, the Dice overlap index of the resulting MECs and LECs was calculated.

In order to cross-validate the results and also to visualize any differences in structural and/or functional whole-brain connectivity between the subregions, tractography analysis was performed by seeding from the rs-fMRI-defined MEC and LEC, and functional connectivity analysis was performed by seeding from the DTI-defined MEC and LEC. The absolute differences between the MEC/LEC structural and functional connectivity maps were also calculated by subtracting the individual maps.

MEC and LEC homologues from all the 20 different segmentation approaches (all ROI combinations, both modalities – see Supplementary Table 2 for an overview) were added together to create a total probability map of MEC vs. LEC predictions based on a combination of DTI and fMRI data. We additionally made a combined-modality MEC vs. LEC segmentation by thresholding and binarizing the probability map.

### 2.6 Signal-to-noise ratio estimation

Signal-to-noise ratio (SNR) measures were estimated within MEC and LEC ROIs obtained from the combined DTI+fMRI segmentations. Temporal SNR (tSNR) of rs-fMRI runs was determined by dividing the mean signal of the ROI by the standard deviation of the mean ROI signal over time. Spatial SNR (sSNR) of rs-fMRI runs was determined by dividing the mean signal of the ROI by the standard deviation of the signal across all voxels in that ROI at each time point, and this was then averaged across all time points in the run. All fMRI SNR estimates were calculated on unsmoothed data, and preprocessed but not denoised data were used for this (because the denoised fMRI data were also de-meaned, which would result in estimated SNR values close to zero). For DTI, a “b0 SNR” was estimated: First, the two first b = 0 images of the acquisition were subtracted from each other, and the standard deviation of the signal difference in the ROI was divided by √2 to obtain a noise estimate of the ROI. Then, the mean signal in the ROI across both b0 images were divided by this noise estimate to obtain the “b0 SNR”. However, note that the effective b-values for “b0” at 3T and 7T were actually b ≈ 5 and b ≈ 60, respectively, and these SNR measures are therefore denoted SNR_b5_ and SNR_b60_.

## 3 Results

Group-averaged and field strength-averaged EC structural connectivity maps for presubiculum+RSC and dCA1pSub+OFC, created from performing tractography on the DTI data, are shown in Figure 1A-B. These results show that the connectivity with presubiculum+RSC is stronger more posteriorly and also slightly medially in the EC, whereas the connectivity with dCA1pSub+OFC is stronger more anteriorly and laterally in the EC. Note that blue color schemes are used throughout for MEC-related connectivity, whereas red color schemes are used for LEC-related connectivity. The resulting DTI-based segmentation of MEC and LEC homologues based on the structural connectivity maps is shown in Figure 1C. There appears to be both a posterior-anterior (PA) and a medial-lateral (ML) orientation of the border between the subregions. The estimated degree of PA orientation of the border for this segmentation was 62.1 ± 0.9 %, whereas the degree of ML orientation was 24.7 ± 5.1 %.

**Figure 1:**
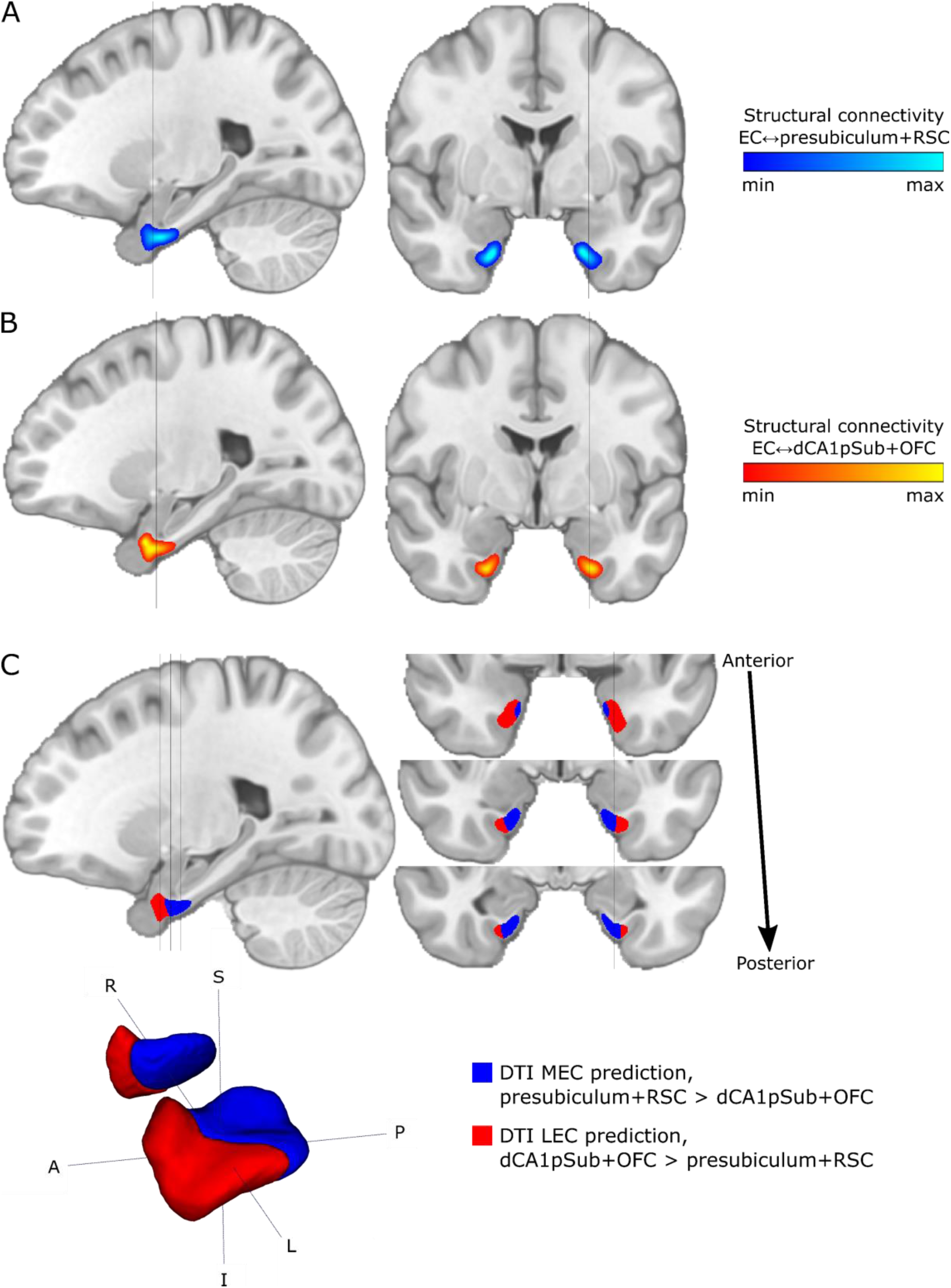
Group-averaged and field strength-averaged structural connectivity maps and MEC vs. LEC segmentations from DTI. Results are shown on selected sagittal (left) and coronal (right) slices in MNI space. Grey lines on sagittal slices show level of adjacent coronal slices, and vice versa. **A:** EC connectivity with presubiculum+RSC, **B:** EC connectivity with dCA1pSub+OFC. **C:** Segmentation of MEC (blue) and LEC (red) homologues based on the connectivity maps, shown both on sagittal and coronal slices and in 3D (bottom row; both hemispheres are shown). S = superior, I = inferior, A = anterior, P = posterior, R = right, L = left.

Corresponding group-averaged and phase encoding direction-averaged EC functional connectivity maps for presubiculum+RSC and dCA1pSub+OFC, created from running functional connectivity analysis on the rs-fMRI data, are shown in Figure 2A-B. Functional connectivity profiles were similar to those from the structural connectivity analysis, where the connectivity with presubiculum+RSC is stronger more posteriorly and medially in the EC, and the connectivity with dCA1pSub+OFC is stronger more anteriorly and slightly laterally in the EC. Note that the structural and functional connectivity maps have a different appearance in the periphery of the maps – this is because of differences between the analysis pipelines: In the DTI analysis, the tractography provided individual EC structural connectivity maps that were already confined to the EC ROI, and these were then co-registered together in the group averaging process. The fMRI analysis, on the other hand, provided whole-brain functional connectivity maps for individual subjects, and these were first co-registered together and then the group-averaged whole-brain connectivity map was masked by an EC ROI. The apparent low connectivity values in the periphery of the structural connectivity maps thus represent registration uncertainties. Nevertheless, the results show a smooth DTI-based segmentation even at the edges of EC, without any apparent misclassification of voxels.

**Figure 2:**
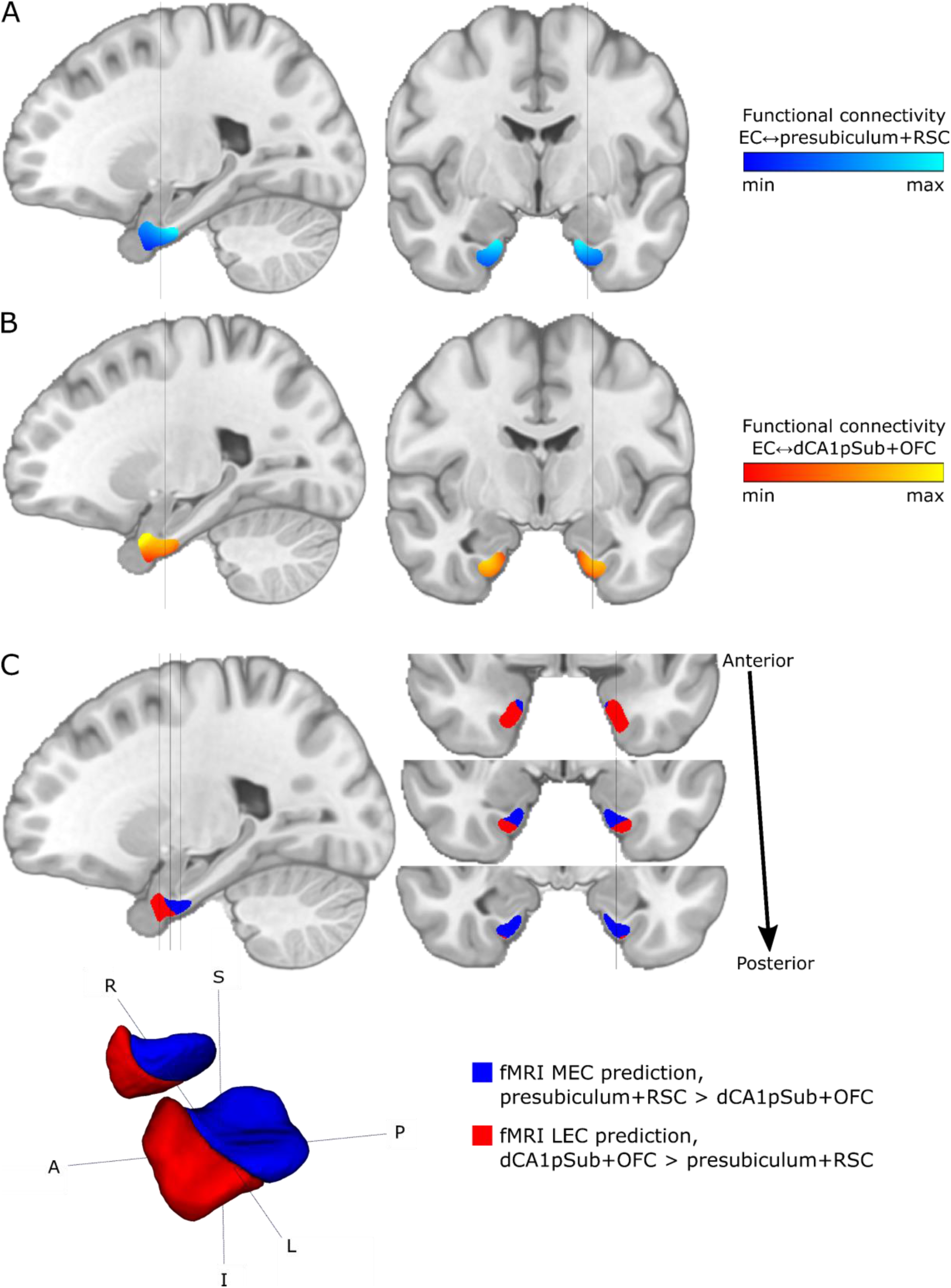
Group-averaged and phase encoding direction-averaged functional connectivity maps and MEC vs. LEC segmentations from rs-fMRI. Results are shown on selected sagittal (left) and coronal (right) slices in MNI space. Grey lines on sagittal slices show level of adjacent coronal slices, and vice versa. **A:** EC connectivity with presubiculum+RSC, **B:** EC connectivity with dCA1pSub+OFC. **C:** Segmentation of MEC (blue) and LEC (red) homologues based on the connectivity maps, shown both on sagittal and coronal slices and in 3D (bottom row; both hemispheres are shown). S = superior, I = inferior, A = anterior, P = posterior, R = right, L = left.

The fMRI-based segmentation of MEC and LEC homologues are shown in Figure 2C, indicating both a PA and an ML orientation of the border between the subregions. However, the ML component appears to be smaller than for the DTI-based segmentation and the PA component relatively larger, with the MEC homologue showing a stronger relative decrease in size in the most anterior coronal slices, and the LEC homologue showing a corresponding relative decrease in size in the most posterior slices. This agrees with the estimated degree of ML orientation of the border for this segmentation of 10.7 ± 3.5 %, and the degree of PA orientation 68.8 ± 2.0 %. Dice overlap index for the DTI- vs. fMRI-based segmentations was 0.87 for both the MEC and the LEC, emphasizing the similarity between the segmentation results from the two different modalities. EC structural and functional connectivity maps, respectively, for the presubiculum, RSC, dCA1pSub and OFC separately are shown in Supplementary Figures 5 and 6. Additional contralateral functional connectivity results, to confirm that the current results are not driven by distance dependencies, are shown in Supplementary Figure 7.

Estimated degrees of PA and ML orientation of the border between the subregions for all 20 segmentation approaches across modalities and seed regions can be found in Supplementary Table 2. In addition, individual segmentation results from both DTI and fMRI analyses are shown in Supplementary Table 3 and Supplementary Figure 8.

What are the resulting functional and structural connectivity networks when seeding from the MEC and LEC homologues as defined from the other imaging modality? Figure 3 shows the functional connectivity networks that result when using the DTI-based MEC and LEC segmentations as seed ROIs in rs-fMRI analysis, while Figure 4 shows the corresponding structural paths that result from using the fMRI-based MEC and LEC segmentations as seed ROIs in DTI tractography. There appears to be a qualitative distinction between the whole-brain resting-state functional connectivity networks of the MEC and LEC. While the MEC whole-brain connectivity maps can be interpreted as associated with components of the canonical default mode network (medial prefrontal cortex, posterior cingulate cortex, angular gyrus)(Raichle, 2015; Reznik et al., 2023), the LEC connectivity shows components which can be recognized to belong to the dorsal attention network (intraparietal sulcus and frontal eye fields, at least in the left hemisphere) (Fox et al., 2006) and salience network (anterior insula, dorsal anterior cingulate cortex (Supplementary Figure 9)) (Menon and Uddin, 2010). The structural connectivity paths seeded from the MEC homologue mainly extend posteriorly along the hippocampal formation passing the presubiculum and towards the RSC. Though the structural connectivity paths seeded from the LEC homologue to some extent also extend posteriorly, these paths additionally take a superior and slightly lateral route passing the dCA1pSub and towards the OFC. Corresponding functional and structural connectivity difference maps of MEC vs LEC are shown in Supplementary Figures 9 and 10, respectively, and a summary of individual cross-validation results is shown in Supplementary Table 4. It is apparent that the MEC and LEC masks defined from fMRI show structural connectivity with the seed regions used in fMRI analysis, and vice versa for the DTI-based segmentation. This data-driven approach therefore serves as a cross-validation of the correspondence between structural and functional connectivity of the EC subregions.

**Figure 3:**
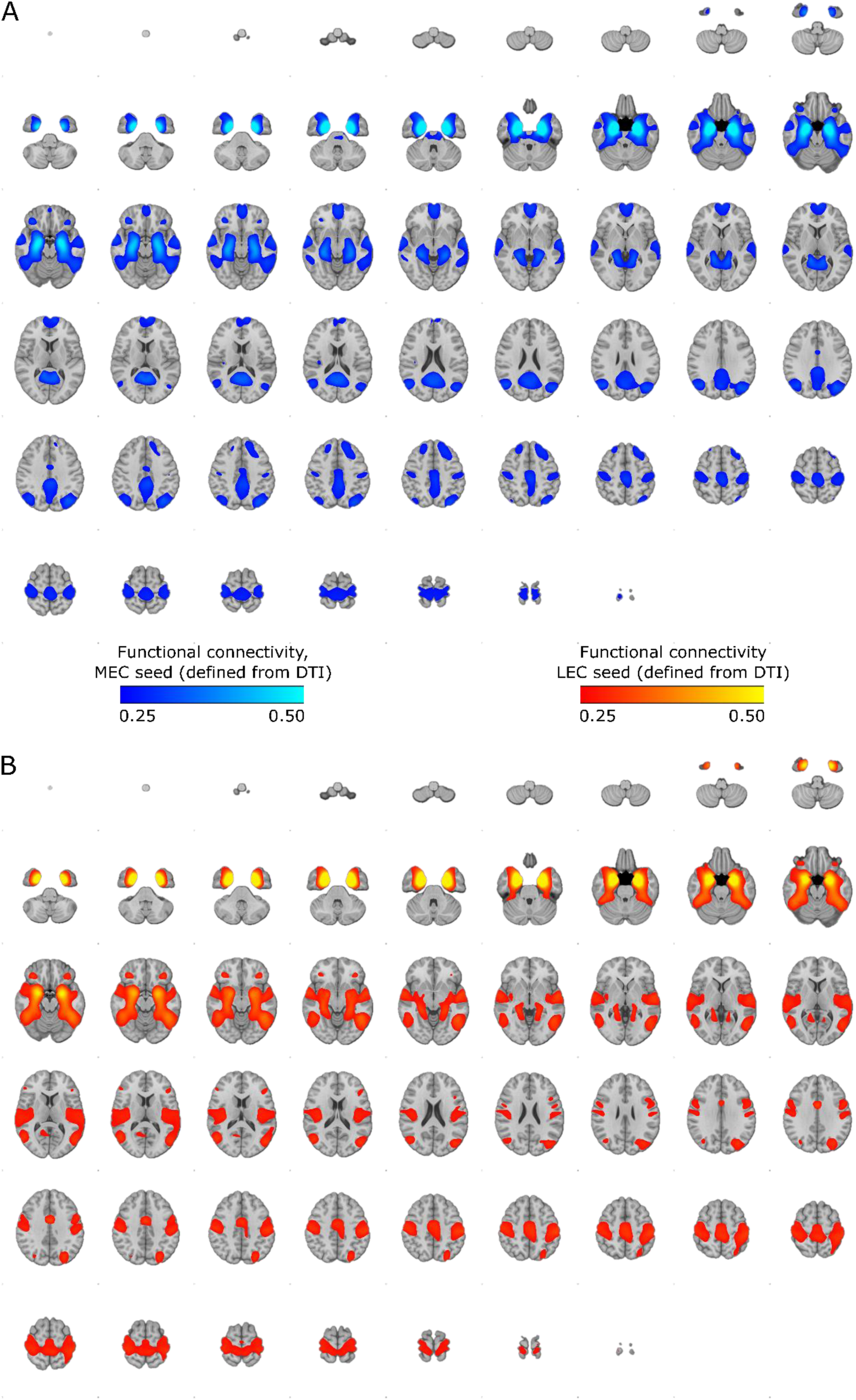
Group-averaged and phase encoding direction-averaged resting-state functional connectivity networks seeded from MEC and LEC ROIs obtained from the DTI-based segmentation. Results are shown on axial slices throughout the brain (MNI space). The connectivity values are group-averaged Fisher Z-transformed Pearson correlation. Whole-brain functional connectivity is shown separately for **A:** MEC seed and **B:** LEC seed. Note that the left side of the images represent the right side of the brain.

**Figure 4:**
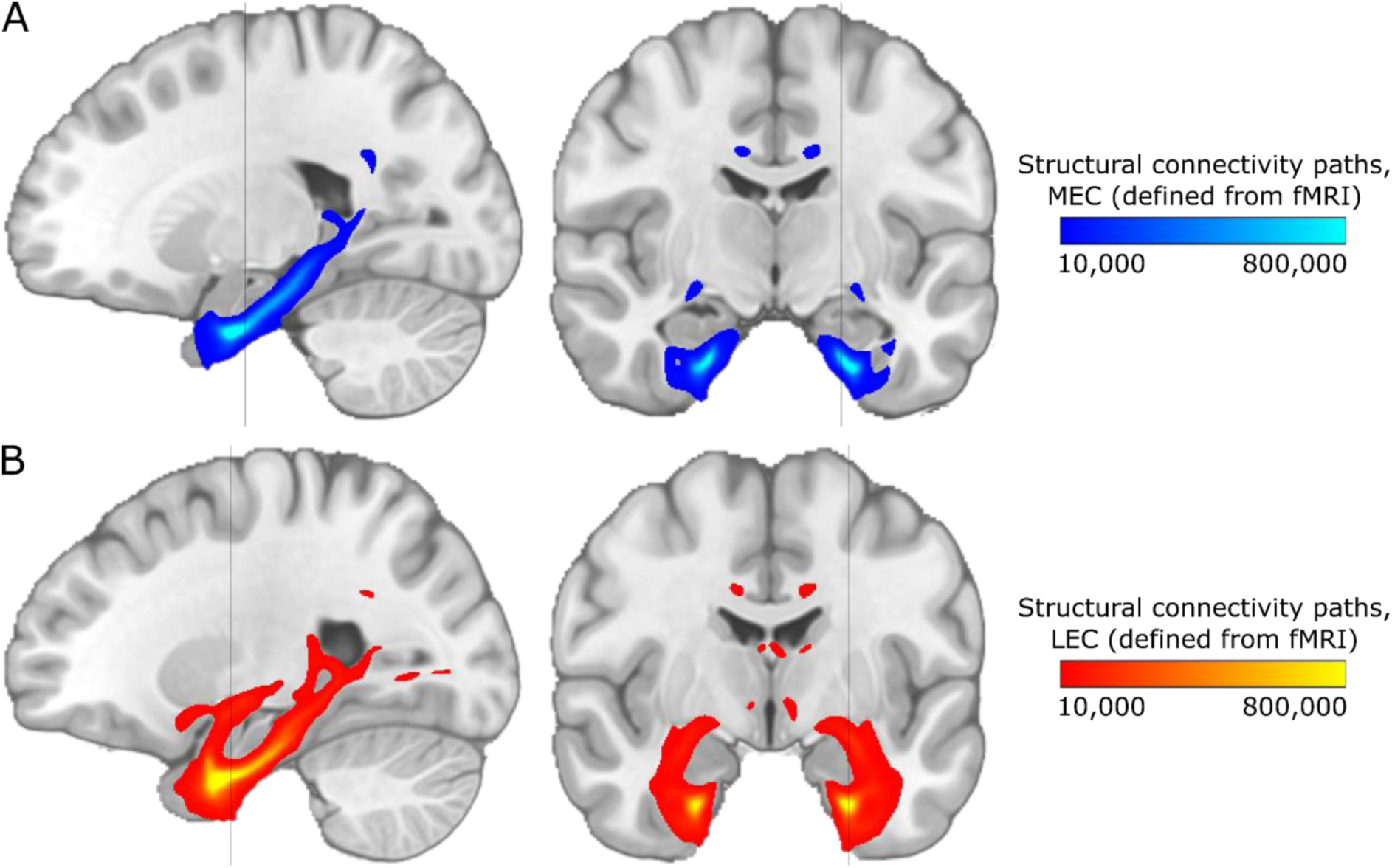
Group-averaged and field strength-averaged structural connectivity paths seeded from MEC and LEC ROIs obtained from the fMRI-based segmentation. Results are shown on selected sagittal (left) and coronal (right) slices in MNI space. Grey line on sagittal slices shows level of adjacent coronal slice, and vice versa. Note that the colormap intensity does not represent the actual number of white matter tracts, but instead scales with the probability that the true path lies in that point. Structural connectivity paths are shown separately for **A:** MEC seed and **B:** LEC seed.

To illustrate the combined results from DTI+MRI, the total probability map of the MEC vs. LEC homologues across the different segmentation approaches is shown in Figure 5A. The sagittal slice shows a relatively sharp transition in probability from MEC to LEC, whereas the coronal slices show a more gradual transition, at least in the most anterior and posterior slices. This reflects a relatively high confidence about the location of the border along the PA axis, but a larger variation concerning the ML component of the border, consistent with the numerical values of PA vs. ML subdivision shown in Supplementary Table 2. Taken together, the maps thus show that the MEC homologue is most likely located in the posteromedial EC (pmEC) based on a combination of seed regions and modalities, whereas the LEC homologue is likely located in the anterolateral EC (alEC). A final combined-modality segmentation of the MEC vs. LEC homologues, based on the probability map in Figure 5A, is shown in Figure 5B. Both the probability map and the final MEC and LEC homologue masks are available in the Supplementary files. As expected, the resulting segmentation looks like a mixture between the DTI- and fMRI-based segmentations in Figure 1C and Figure 2C, respectively. The estimated degree of PA orientation of the border for the DTI+fMRI segmentation was 65.5 ± 2.3 %, while the degree of ML orientation was 18.9 ± 4.5 %. This combined-modality approach thus provides high confidence that the pmEC represents the human homologue of MEC, whereas the alEC represents the LEC homologue.

**Figure 5:**
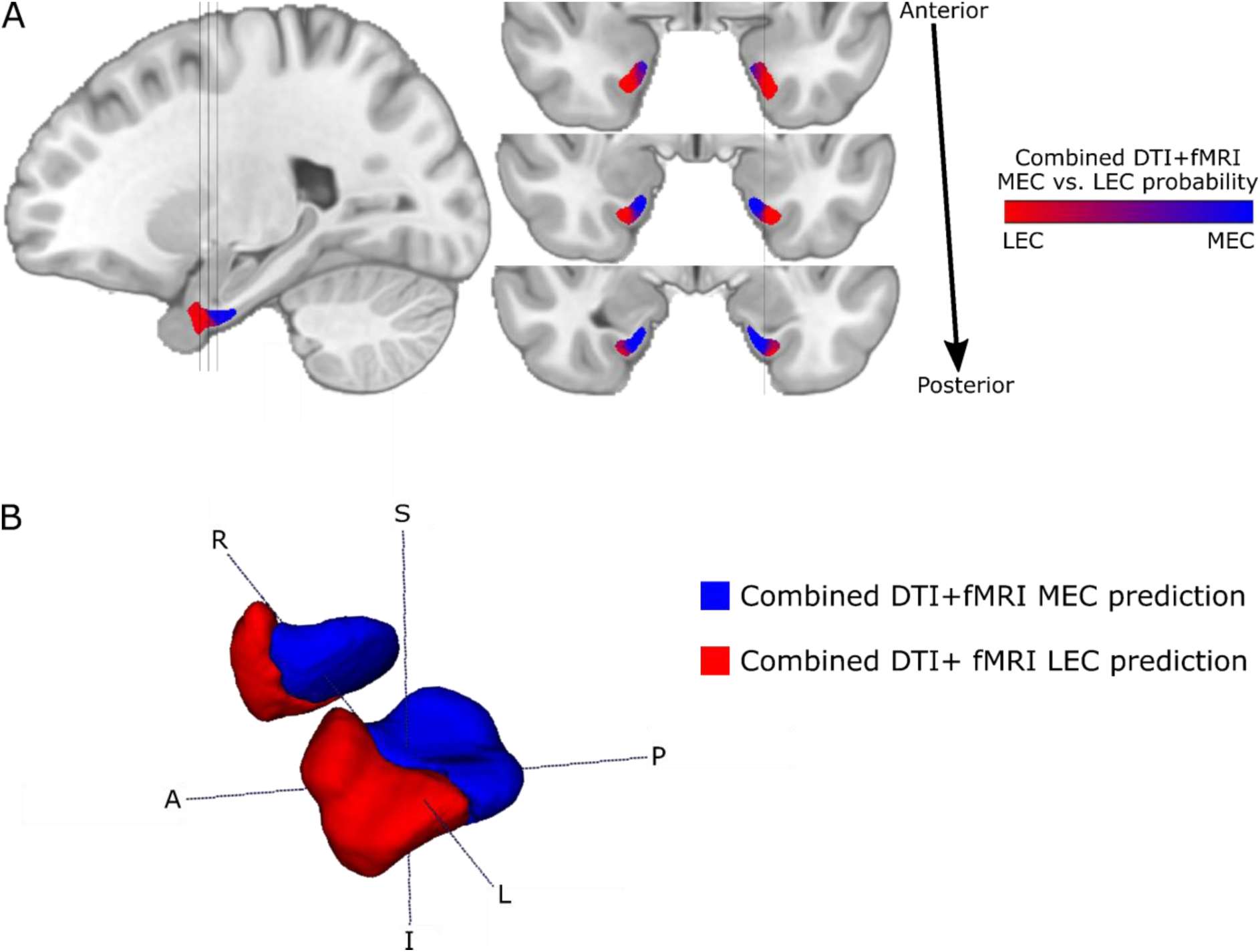
Combined DTI+fMRI results. **A**: Total combined probability of MEC vs. LEC homologue predictions across all the 20 segmentation approaches (see Supplementary Table 2) from both DTI and fMRI. Results are shown on selected sagittal (left) and coronal (right) slices in MNI space. Grey lines on sagittal slice show level of adjacent coronal slices, and vice versa. **B:** Combined DTI+fMRI segmentation of MEC vs. LEC based on the combined probability map. Results from both hemispheres are shown in 3D. S = superior, I = inferior, A = anterior, P = posterior, R = right, L = left.

SNR estimates in MEC and LEC ROIs obtained from the combined DTI+fMRI segmentation are shown in Table 1. For the fMRI data, both tSNR and sSNR are significantly higher in MEC than in LEC for the PA phase encoding direction, and significantly higher in LEC than in MEC for the AP phase encoding direction. For the DTI data, SNR is significantly higher in LEC than in MEC at 3T, whereas there is no significant difference between MEC and LEC at 7T. While sSNR is generally higher for the PA phase encoding direction than for AP, there are no obvious differences for tSNR across phase encoding directions. For DTI, SNR is generally higher for 3T than for 7T.

**Table 1:** SNR measures in MEC and LEC ROIs defined from the combined DTI+fMRI segmentation. Numbers are given as mean ± standard deviation across participants, separately for posterior-anterior (PA) and anterior-posterior (AP) phase encoding directions for fMRI data, and separately for 3T and 7T field strengths for DTI data. SNR comparisons between MEC vs. LEC marked with an asterisk (*) are statistically significant (p < 0.05).

|  |  |  | MEC | LEC |
| --- | --- | --- | --- | --- |
| fMRI data | tSNR | PA* | 84 ± 31 | 77 ± 29 |
| | | AP* | 65 ± 27 | 100 ± 36 |
| | sSNR | PA* | 2.64 ± 1.10 | 2.50 ± 1.04 |
| | | AP* | 1.78 ± 0.64 | 2.34 ± 0.99 |
| DTI data | SNR<sub>b5</sub> | 3T* | 14.8 ± 3.0 | 16.5 ± 2.4 |
| | SNR<sub>b60</sub> | 7T | 11.3 ± 2.1 | 11.3 ± 2.5 |

## 4 Discussion

In this study, we used both DTI and rs-fMRI data from 103 healthy adults to investigate structural and functional connectivity of the human EC, aiming to predict the locations of the human homologues of MEC and LEC as defined in experimental animal studies based on cytoarchitectonics and connectivity criteria. We found that using either DTI or rs-fMRI to segment EC subregions predicted approximately the same two parts, and the combinatorial usage resulted in a posteromedial (pmEC) an anterolateral (alEC) part with a rather confined border region between them. The connectivity profiles of these two parts align with animal connectivity data, such that human pmEC matches MEC, while human alEC matches LEC. Interestingly, while pmEC showed increased connectivity preferentially with the default mode network, the alEC exhibited increased connectivity with regions in the dorsal attention and salience networks. Our findings support the general subdivision of the EC into pmEC and alEC, as suggested in previous studies (Maass et al., 2015; Navarro Schröder et al., 2015; Syversen et al., 2021), and allow us to define a border region between the two with high certainty.

The modalities yielded similar subdivisions along the posterior-anterior axis but resulted in some variation along the medial-lateral axis. While previous fMRI studies reported a subdivision primarily along the PA axis, our previous DTI study found a somewhat lesser degree of PA orientation and higher degree of ML orientation of the border between the subregions than the fMRI studies, although note that the PA component still remained larger than the ML component (Maass et al., 2015; Navarro Schröder et al., 2015; Syversen et al., 2021). This is consistent with our current findings, where both the DTI- and fMRI-based final segmentations show a larger degree of PA orientation of the border than what is the case for the ML orientation, and fMRI yields more of a PA orientation than DTI. The final combined-modality segmentation yields a substantial degree of both PA and ML orientation, although still with a larger PA than ML component. The degrees of PA and ML orientation of the border in our previous DTI study using the same seed regions for segmentation were 58% and 19%, respectively (Syversen et al., 2021), while the results of the current study were 62% and 25%. This suggests that the DTI results are fairly reproducible across cohorts and MRI acquisition protocols. For fMRI, there are no earlier studies that can be used as a direct comparison, although the alEC and pmEC segmentations from two previous fMRI studies had a higher degree of PA subdivision and a lower degree of ML subdivision than our fMRI results (Supplementary Table 2, Supplementary Figure 11) (Maass et al., 2015; Navarro Schröder et al., 2015). However, the latter studies used different seed regions than the current study.

Although using structural and functional connectivity approaches resulted in qualitatively similar subdivisions of the EC, it is uncertain whether the abovementioned variations in the PA vs. ML orientation of the border between pmEC and alEC stem from real differences in structural vs. functional connectivity, or if they are caused by inherent differences or limitations in the two imaging modalities and/or subsequent analyses. While structural connectivity constitutes the framework for functional connectivity, functionally connected regions are not necessarily directly (monosynaptically) structurally connected, and vice versa (Rykhlevskaia et al., 2008). Irrespective of this potential variation, tractography and functional connectivity analyses performed on pmEC and alEC ROIs defined through a cross-validation approach from the opposite modality showed differential patterns of connections that were similar (and thus generalizable) across functional and structural connectivity. This is in line with other studies where both types of connectivity have been investigated and compared in the hippocampal memory system in the same subjects (Huang et al., 2021; Ma et al., 2022; Rolls et al., 2022), although the functional network maps here (Figure 3) show more extensive distant connections than the structural path maps (Figure 4). However, note that the appearance of these maps depends on the selected thresholding level, and that structural and functional connectivity analysis have inherently different distance dependencies. Differences in both the analysis methods and data acquisition between DTI and fMRI is a possible source of variation between structural and functional connectivity results. The fMRI data has a coarser resolution, potentially making it less sensitive to connectivity differences along the narrow medial-lateral axis of the EC. In addition, the DTI data is acquired using a spin echo MRI sequence whereas the fMRI data is acquired using gradient echo, which can further contribute to differences in data quality and artefact levels.

In rodents, there is a sharply defined border between MEC and LEC based on both cytoarchitecture and differential anatomical projections (Kjonigsen et al., 2011; Witter, 2007). In non-human primates, however, the topography of projections in the EC does not appear to adhere to any distinct cytoarchitectonic division, but instead shows a gradient along the posteromedial to anterolateral axis (Witter and Amaral, 2021). One might therefore expect a similar topographical gradient in humans, as shown in the present and previous studies (Maass et al., 2015; Navarro Schröder et al., 2015; Syversen et al., 2021). Thus, the question about whether there is a sharp or a gradual border between the human pmEC and alEC has not yet been clearly answered. We do however demonstrate that combining segmentation approaches using different modalities and combinations of seed regions results in a probability map with a gradual (although relatively confined) increase and decrease in confidence of pmEC and alEC locations along the posteromedial and anterolateral axes, respectively. Note however, that our results could be influenced by the low number of seed regions used. Connectivity from the EC to even more brain regions should be investigated in the future to more accurately map the topography of connections, as different regions might be structurally and/or functionally connected with distinct subparts of MEC and LEC (Grande et al., 2022; Reznik et al., 2023; Witter and Amaral, 2021; Witter et al., 2017). Cytoarchitectonic analyses support the subdivision of human EC into multiple subregions (Insausti et al., 1995; Krimer et al., 1997; Oltmer et al., 2022), however, these cannot be directly compared to our MRI data since that would require a level of resolution and data quality beyond what has currently been possible with in vivo MRI of the MTL. This will hopefully continue to improve with the advent of new imaging approaches, technologies, and analyses (Dalton et al., 2022; Reznik et al., 2023). Another future possibility may be to use a qualitative comparison (van Oort et al., 2018) and to add histological validation.

Equal sizes of MEC and LEC were assumed in our segmentation process, based on rodent data (Merrill et al., 2000). This might not be true for humans, but this assumption was made due to lack of evidence of what the real relative sizes of the subregions should be. One of the previous studies segmenting the EC into alEC and pmEC also assumed equal sizes (Navarro Schröder et al., 2015), while in another study the resulting alEC was larger than the pmEC (Maass et al., 2015). Some might argue that this is reasonable, as the LEC has a more elaborate connectivity to other brain areas than the MEC, at least in rodents (Bota et al., 2015; Doan et al., 2019). Nevertheless, our analysis methods are not suitable to confidently determine the relative sizes of human MEC vs. LEC, as both the structural and functional connectivity values estimated are relative numbers that depend on a number of variables.

Our study has some limitations. First, the ROIs used as seed regions for the analyses are obtained from automated parcellation and then registered to each participants’ brain. Although manual adjustments to the MNI space ROIs were made and all ROIs were also masked by the participants’ individually parcellated ROIs, remaining anatomical inaccuracies in ROIs could have affected the results. For example, there are individual differences in the shape of the EC and the depth of the collateral sulcus, which makes up the lateral border of the EC (Insausti et al., 1998; Oltmer et al., 2023). This is a general limitation of group studies compared with e.g. individual precision imaging of the human MTL (Reznik et al., 2023). On a similar note, the structural and functional connectivity analyses were performed in separate spaces (native vs. MNI, respectively), which introduces some uncertainty to the comparison between them – but this was how the preprocessed data were provided from the HCP. Another limitation is that the EPI sequences are not particularly optimized for the MTL, and this region is therefore subject to relatively low SNR, geometric distortions and signal dropouts in the images, especially at 7T. To increase the sensitivity of detecting functional correlations in this region, the fMRI data were smoothed with a Gaussian kernel. However, this reduces the spatial resolution and anatomical specificity of the results. For both DTI and rs-fMRI, the results could be influenced by the fact that some of the seed regions (presubiculum and dCA1pSub) are located relatively close to the EC, although this was attempted to be mediated by also choosing distant, neocortical seed regions (RSC and OFC). Contralateral fMRI analyses (Supplementary Figure 7) nevertheless indicate that our fMRI results are not strongly driven by distance dependencies. At last, there are differences in SNR for the MEC and LEC segmentations. Nevertheless, these differences are fairly balanced – at least for the fMRI data – as the MEC/LEC SNR ratios for PA phase encoding direction are practically inversed for AP, and our final segmentations are averaged across both phase encoding directions for fMRI and both field strengths for DTI.

## 5 Conclusions

Our study provides converging evidence that both DTI and rs-fMRI yield similar subdivision of the human EC into posteromedial (pmEC) and anterolateral (alEC) parts. This extends and bridges findings from previous studies where DTI and fMRI were separately investigated, and more clearly reveals a confined border between pmEC and alEC. Furthermore, the resulting subregions show characteristic patterns of connections that are similar across functional and structural connectivity, with a differential connectivity profile relating to the default mode network (pmEC) and to the dorsal attention and salience networks (alEC), respectively. Future studies should map the human EC connectivity to even more brain regions and investigate connections to distinct subregions, to better understand their topography. The pmEC and alEC as likely homologues of MEC and LEC, when applied to other cognitive and translational MRI studies, will strongly facilitate the possibilities to compare results between studies and between species.

## Supplementary materials

- **Supplementary figures and tables:** Supplementary Figures 1-11, Supplementary (supplementary_figures_tables.docx).
- **Supplementary files:** MEC and LEC homologue masks in MNI152-09b space, defined from the combined-modality segmentation shown in Figure 5B, as well as the MEC vs. LEC probability map shown in Figure 5A (MEC_LEC_segmentations.zip).

## Acknowledgements

The authors would like to thank Pål Erik Goa for assistance on MRI physics and SNR questions, and for detailed discussion and general support.

Data were provided by the Human Connectome Project, WU-Minn Consortium (Principal Investigators: David Van Essen and Kamil Ugurbil; 1U54MH091657) funded by the 16 NIH Institutes and Centers that support the NIH Blueprint for Neuroscience Research; and by the McDonnell Center for Systems Neuroscience at Washington University.

This study was supported by a PhD grant from the Faculty of Medicine and Health Sciences at NTNU awarded to TNS. CFD’s research is further supported by the Max Planck Society, the European Research Council (ERC-CoG GEOCOG 724836), the Kavli Foundation, the Jebsen Foundation, Helse Midt Norge and The Research Council of Norway (223262/F50; 197467/F50).

The funding sources had no role in study design, data collection, data analysis, data interpretation or writing of the report.

---

**Supplementary Figure 1:**
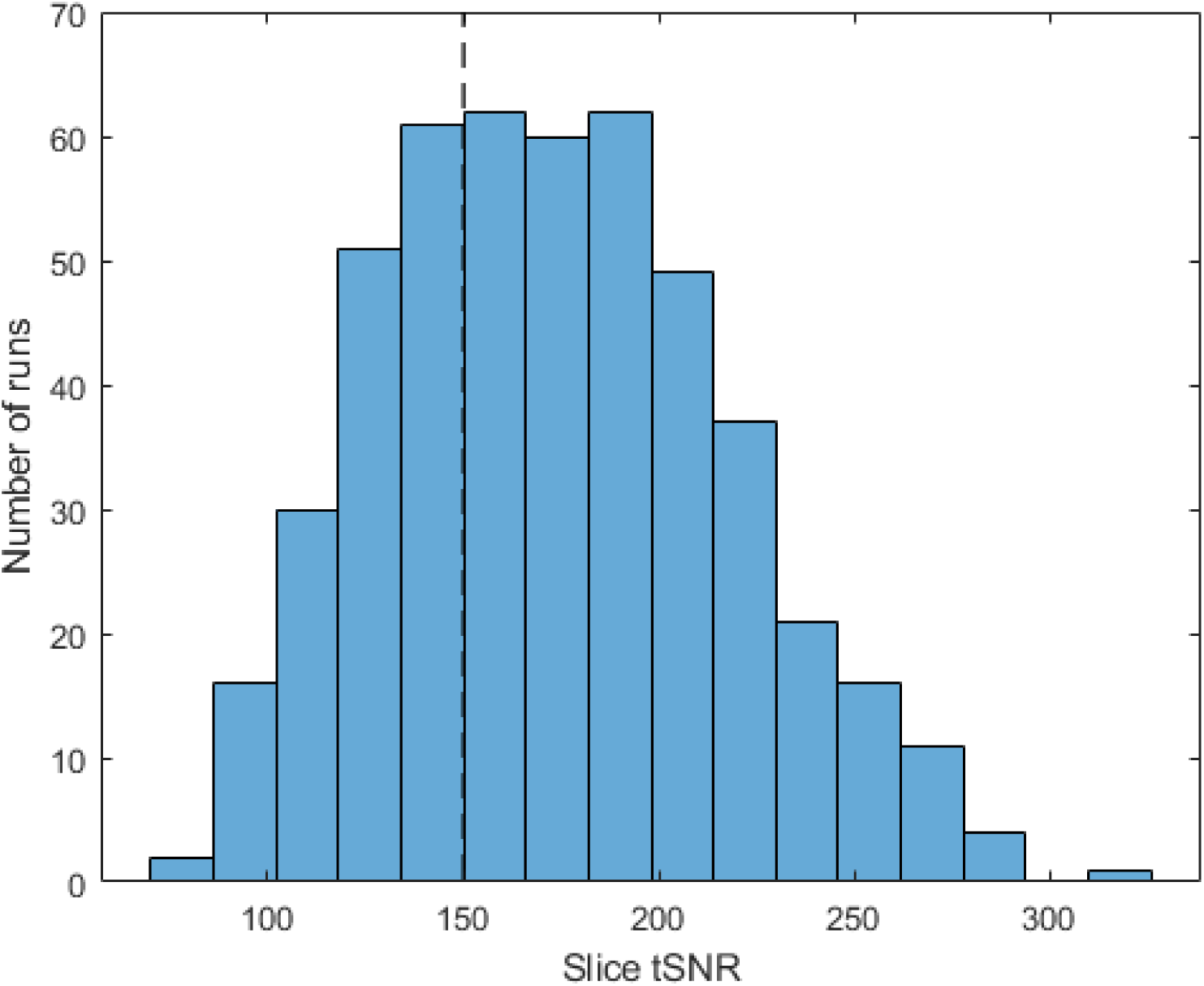
Histogram showing the whole-brain tSNR for quality control before exclusion of rs-fMRI data. fMRI runs with slice tSNR ≤ 150 were excluded (shown by dashed line). In total, 30 participants were excluded due to insufficient slice tSNR, and 160 out of 483 fMRI runs (33%) were excluded. The mean slice tSNR for all runs was 173, and the mean slice tSNR for all included runs was 197.

**Supplementary Figure 2:**
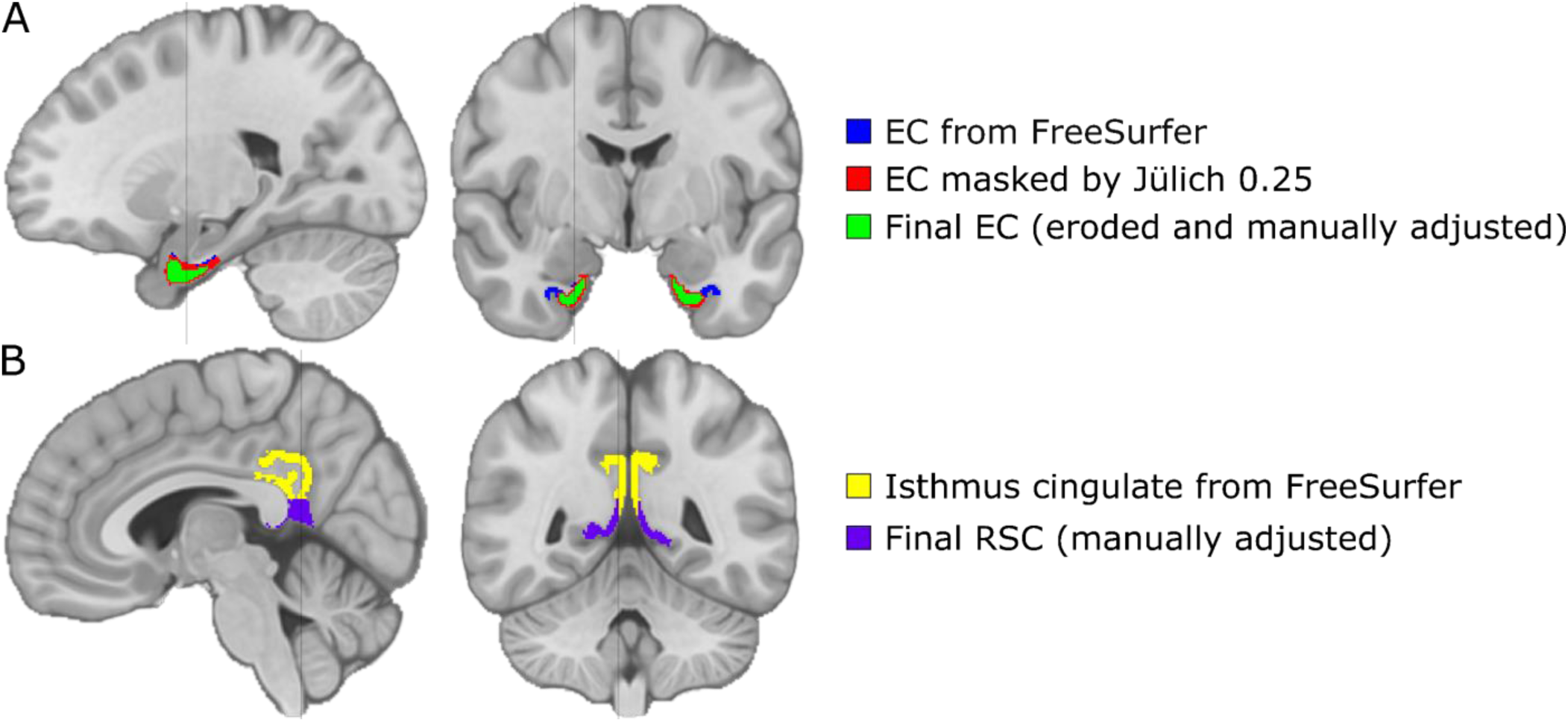
Adjustments of EC and RSC ROIs. ROIs are shown on a representative sagittal (left) and a coronal (right) slice in MNI space. Grey line on sagittal slices shows level of adjacent coronal slice, and vice versa. **A:** EC ROI from FreeSurfer is shown in blue, this was masked by a Jülich Atlas EC ROI thresholded at 0.25 to create the red ROI. It was then eroded and manually adjusted to obtain the final EC ROI shown in green. **B:** Isthmus cingulate ROI from FreeSurfer is shown in yellow, this was manually tailored to obtain the final RSC ROI shown in purple.

**Supplementary Figure 3:**
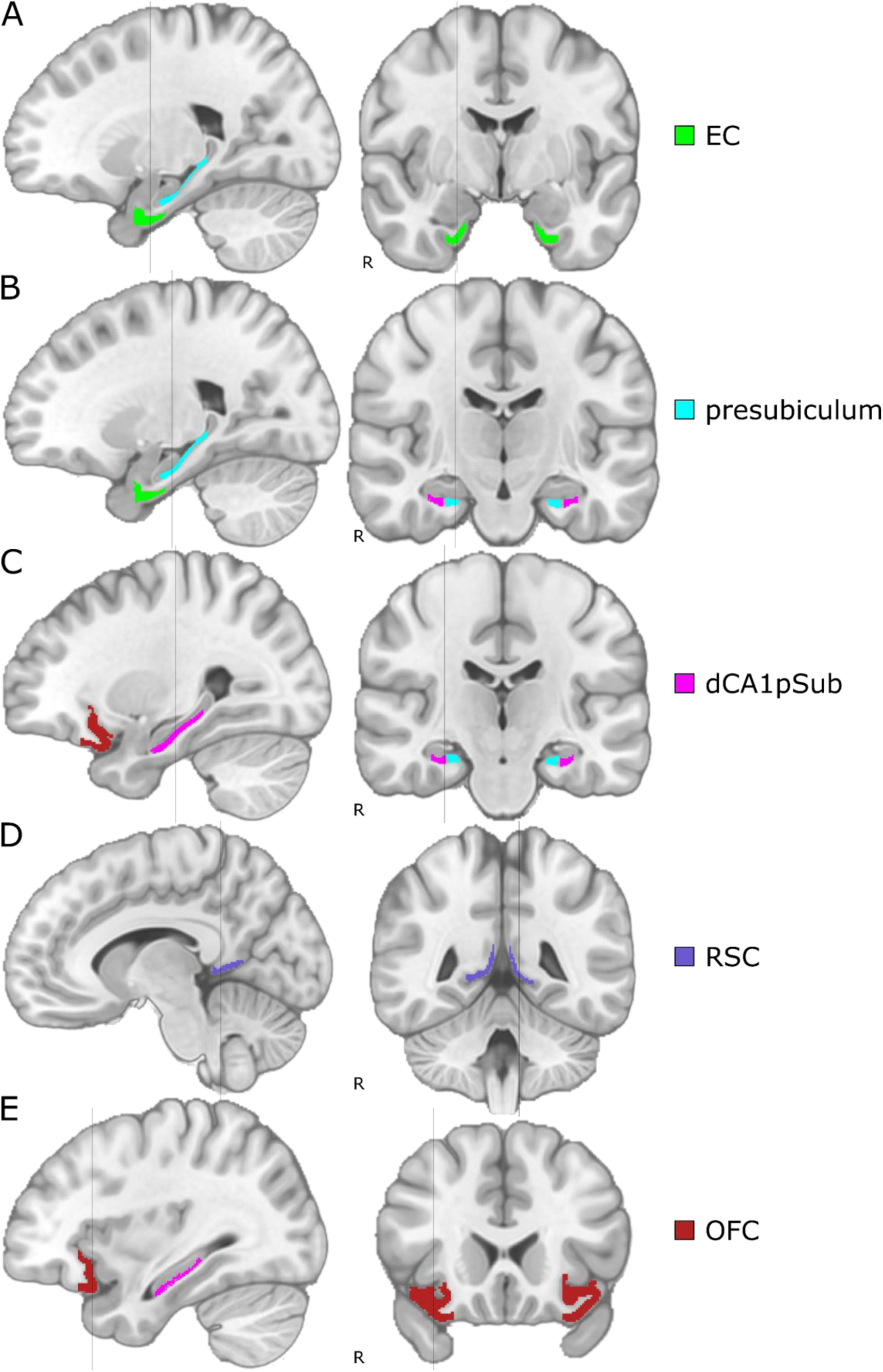
ROIs used for DTI and rs-fMRI analysis. All ROIs are shown on a representative sagittal (left) and a coronal (right) slice in MNI space, with “R” denoting the right side of the brain. Grey line on sagittal slices shows level of adjacent coronal slice, and vice versa. **A:** EC (green), **B:** presubiculum (light blue), **C:** dCA1pSub (pink), **D:** RSC (purple), **E:** OFC (dark red).

**Supplementary Table 1:** Sizes of segmented MEC and LEC, and resulting MEC/LEC size ratio, from the group-level segmentation process before scaling of connectivity maps. Especially for fMRI analyses, values of EC connectivity maps from close (presubiculum and dCA1pSub) versus distant (RSC and OFC) ROIs were so different that the segmented MEC or LEC would comprise the whole EC volume for some segmentation approaches. Also shown are the range of values (min–max) of the connectivity maps used for the segmentations.

| Segmentation |  | Size (# voxels) |  | MEC/LEC<br>size ratio | Connectivity map range |  |
| --- | --- | --- | --- | --- | --- | --- |
|  |  | MEC | LEC |  | MEC seed | LEC seed |
| DTI | Presubiculum/dCA1pSub | 13668 | 11523 | 1.19 | 0.00–0.20 | 0.00–0.18 |
|  | RSC/OFC | 22127 | 5031 | 4.40 | 0.00–0.23 | 0.00–0.20 |
|  | Presubiculum/OFC | 16231 | 8047 | 2.02 | 0.00–0.20 | 0.00–0.20 |
|  | RSC/dCA1pSub | 17740 | 8201 | 2.16 | 0.00–0.23 | 0.00–0.18 |
|  | Presubiculum+RSC/dCA1pSub+OFC | 17815 | 7935 | 2.25 | 0.00–0.20 | 0.00–0.14 |
| fMRI | Presubiculum/dCA1pSub | 5294 | 20456 | 0.26 | 0.19–0.74 | 0.20–0.68 |
|  | RSC/OFC | 18029 | 7721 | 2.34 | 0.12–0.42 | 0.11–0.34 |
|  | Presubiculum/OFC | 25750 | 0 | ∞ | 0.19–0.74 | 0.11–0.34 |
|  | RSC/dCA1pSub | 0 | 25750 | 0 | 0.12–0.42 | 0.20–0.68 |
|  | Presubiculum+RSC/dCA1pSub+OFC | 16856 | 8894 | 1.90 | 0.15–0.53 | 0.14–0.40 |

**Supplementary Table 2:** Degree of posterior-anterior (PA) and medial-lateral (ML) orientation of the border between MEC and LEC for the 20 different segmentation approaches. For DTI, segmentation was performed separately for 3T and 7T field strengths, while for fMRI, segmentation was performed separately for posterior-anterior (PA) and anterior-posterior (AP) phase encoding directions. In addition, PA and ML subdivision from previous fMRI studies (Navarro Schröder et al. eLife 2015, Maass et al. eLife 2015) have also been calculated for comparison. The degree of PA and ML orientation is given as a percentage between 0 and 100%, dependent on the angle between the MEC-LEC center of gravity vector and the pure PA or ML vector, respectively. All numbers are given as the mean of both hemispheres ± mean absolute deviation.

| Segmentation approach |  |  | Posterior-anterior (PA) axis |  | Medial-lateral (ML) axis |  |
| --- | --- | --- | --- | --- | --- | --- |
|  |  |  | Angle (°) | % PA | Angle (°) | % ML |
| DTI | 3T | Presubiculum/dCA1pSub | 45.4 ± 3.5 | 49.6 ± 3.9 | 29.0 ± 7.9 | 67.8 ± 8.7 |
|  |  | RSC/OFC | 35.2 ± 5.4 | 60.8 ± 6.0 | 86.3 ± 7.6 | 4.2 ± 8.4 |
|  |  | Presubiculum/OFC | 39.5 ± 0.6 | 56.1 ± 0.7 | 54.6 ± 0.2 | 39.3 ± 0.3 |
|  |  | RSC/dCA1pSub | 27.2 ± 0.5 | 69.7 ± 0.6 | 78.7 ± 5.2 | 12.6 ± 5.8 |
|  |  | Presubiculum+RSC/dCA1pSub+OFC | 30.8 ± 0.1 | 65.7 ± 0.1 | 69.0 ± 2.4 | 23.3 ± 2.7 |
|  | 7T | Presubiculum/dCA1pSub | 47.9 ± 5.6 | 46.7 ± 6.2 | 30.8 ± 10.2 | 65.7 ± 11.3 |
|  |  | RSC/OFC | 44.2 ± 5.8 | 50.9 ± 6.5 | 82.4 ± 1.3 | 8.4 ± 1.5 |
|  |  | Presubiculum/OFC | 53.6 ± 0.5 | 40.5 ± 0.5 | 46.8 ± 6.3 | 48.0 ± 7.0 |
|  |  | RSC/dCA1pSub | 33.8 ± 3.4 | 62.5 ± 3.8 | 78.2 ± 7.5 | 13.1 ± 8.3 |
|  |  | Presubiculum+RSC/dCA1pSub+OFC | 39.1 ± 1.7 | 56.6 ± 1.9 | 65.8 ± 7.1 | 26.9 ± 7.9 |
| fMRI | PA | Presubiculum/dCA1pSub | 44.9 ± 3.8 | 50.1 ± 4.2 | 33.5 ± 5.6 | 62.7 ± 6.2 |
|  |  | RSC/OFC | 29.0 ± 2.4 | 67.8 ± 2.7 | 81.1 ± 4.0 | 9.9 ± 4.4 |
|  |  | Presubiculum/OFC | 28.8 ± 1.8 | 68.0 ± 2.0 | 82.8 ± 2.8 | 8.0 ± 3.1 |
|  |  | RSC/dCA1pSub | 79.4 ± 6.6 | 11.8 ± 7.4 | 18.6 ± 8.1 | 79.3 ± 9.0 |
|  |  | Presubiculum+RSC/dCA1pSub+OFC | 28.7 ± 2.0 | 68.1 ± 2.2 | 79.3 ± 3.6 | 11.9 ± 4.0 |
|  | AP | Presubiculum/dCA1pSub | 34.1 ± 0.9 | 62.1 ± 0.9 | 60.2 ± 2.4 | 33.1 ± 2.7 |
|  |  | RSC/OFC | 28.5 ± 2.5 | 68.4 ± 2.7 | 84.4 ± 3.1 | 6.2 ± 3.4 |
|  |  | Presubiculum/OFC | 19.5 ± 0.6 | 78.3 ± 0.7 | 77.9 ± 2.9 | 13.5 ± 3.2 |
|  |  | RSC/dCA1pSub | 42.4 ± 6.0 | 52.9 ± 6.7 | 79.7 ± 3.5 | 11.4 ± 3.9 |
|  |  | Presubiculum+RSC/dCA1pSub+OFC | 27.1 ± 1.9 | 69.9 ± 2.1 | 81.7 ± 2.8 | 9.3 ± 3.1 |
| Navarro Schröder et al. eLife 2015 (fMRI) |  |  | 6.8 ± 1.8 | 92.4 ± 2.0 | 85.5 ± 0.7 | 5.0 ± 0.8 |
| Maass et al. eLife 2015 (fMRI) |  |  | 6.6 ± 0.5 | 92.7 ± 0.6 | 87.9 ± 0.9 | 2.4 ± 1.0 |

**Supplementary Figure 4:**
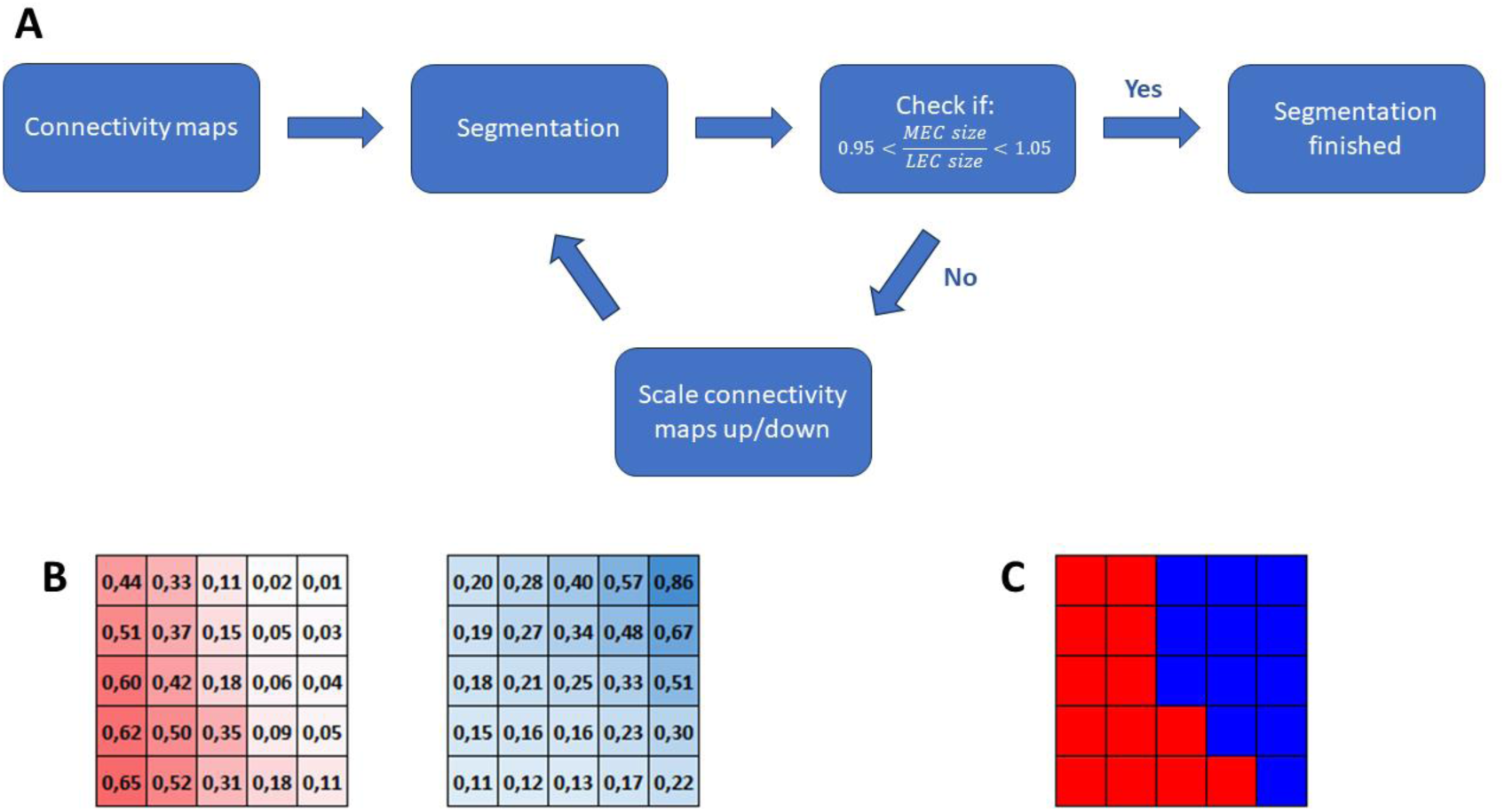
Illustration of the segmentation and scaling process. Part **A** of the figure shows the scheme of the iterative scaling of the segmentations. The raw connectivity maps from DTI and fMRI analysis are used for the segmentation, by comparing the maps pixel by pixel and determining for each pixel which connectivity map has the strongest value. As an example, if the red and blue connectivity maps in **B** are compared, and in the upper left pixel the values are red=0.44 and blue=0.20, that pixel is classified as “red” (see **C**). After the segmentation, the sizes of the resulting MEC and LEC are compared. If the MEC/LEC size ratio is between 0.95 and 1.05, the segmentation is approved and the whole process is finished. If the size ratio is outside this range, however, the connectivity maps are scaled – meaning that one of the maps is multiplied by a small factor. After this, the segmentation step is performed again on the scaled connectivity maps, and the new MEC/LEC sizes are calculated. This circle will continue until the size ratio is within the approved range.

**Supplementary Figure 5:**
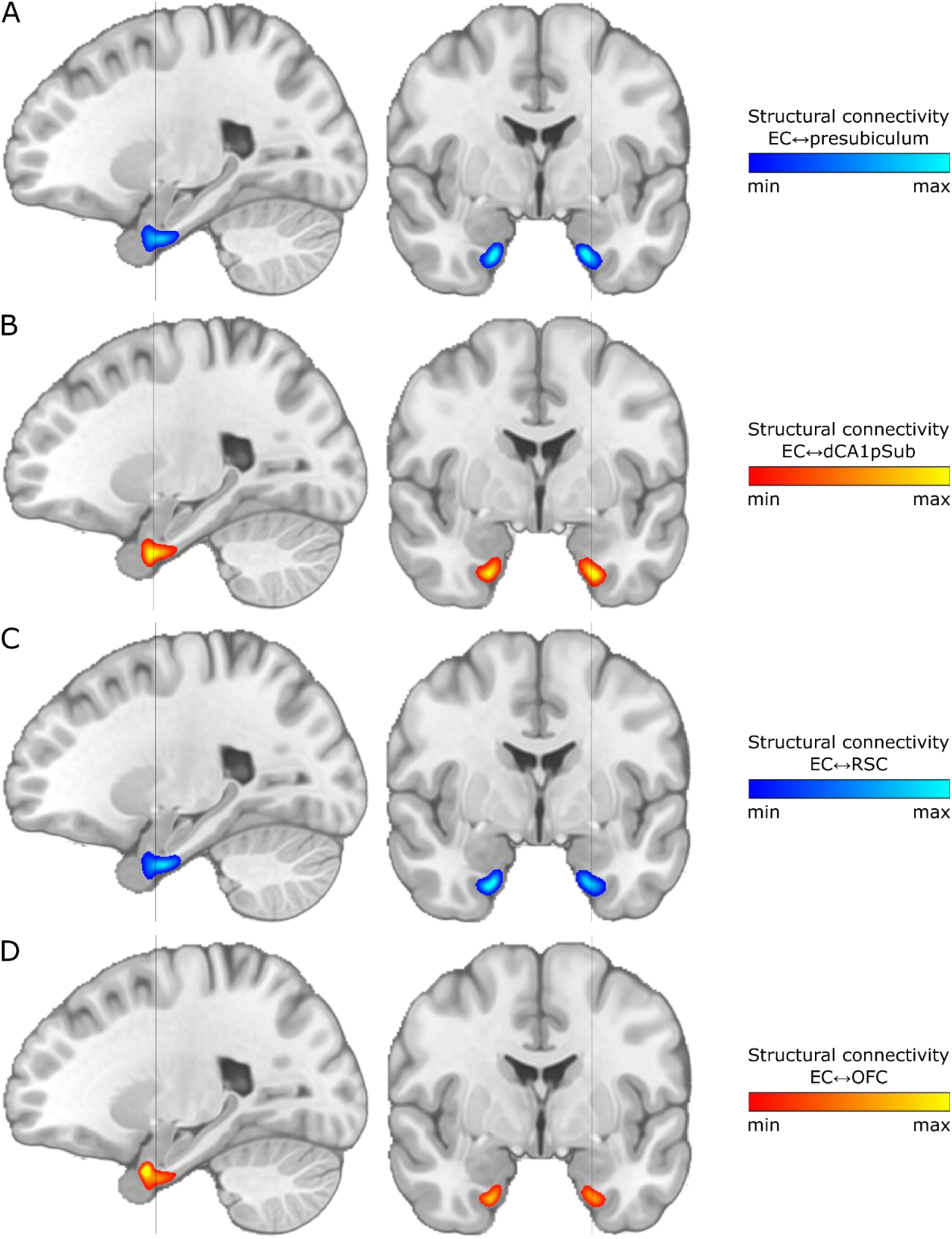
Group-averaged and field strength-averaged structural connectivity maps from DTI. Results are shown on selected sagittal (left) and coronal (right) slices in MNI space. Grey line on sagittal slices shows level of adjacent coronal slice, and vice versa. **A:** EC connectivity with presubiculum, **B:** EC connectivity with dCA1pSub, **C:** EC connectivity with RSC, **D:** EC connectivity with OFC.

**Supplementary Figure 6:**
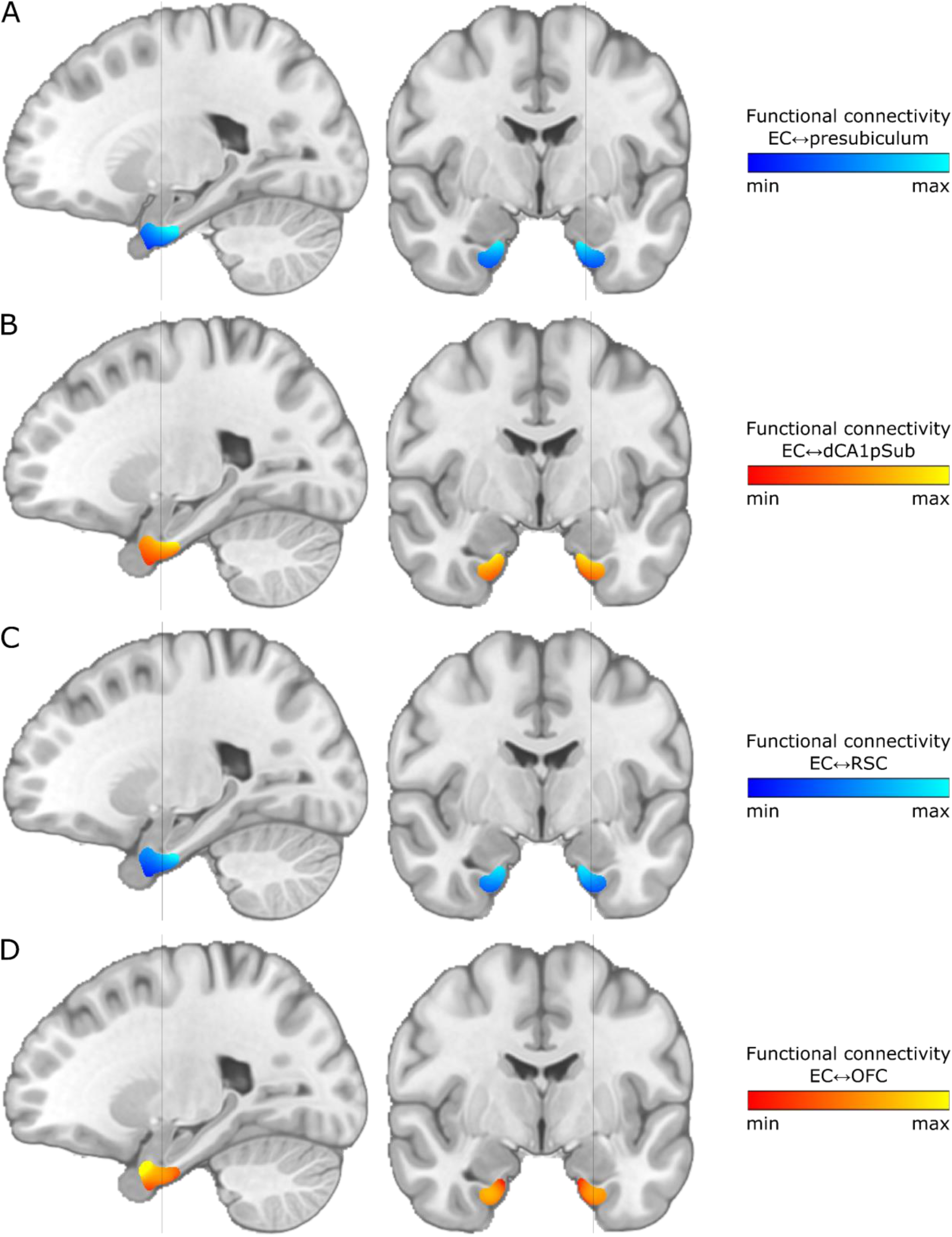
Group-averaged and phase encoding direction-averaged functional connectivity maps from rs-fMRI. Results are shown on selected sagittal (left) and coronal (right) slices in MNI space. Grey line on sagittal slices shows level of adjacent coronal slice, and vice versa. **A:** EC connectivity with presubiculum, **B:** EC connectivity with dCA1pSub, **C:** EC connectivity with RSC, **D:** EC connectivity with OFC.

**Supplementary Figure 7:**
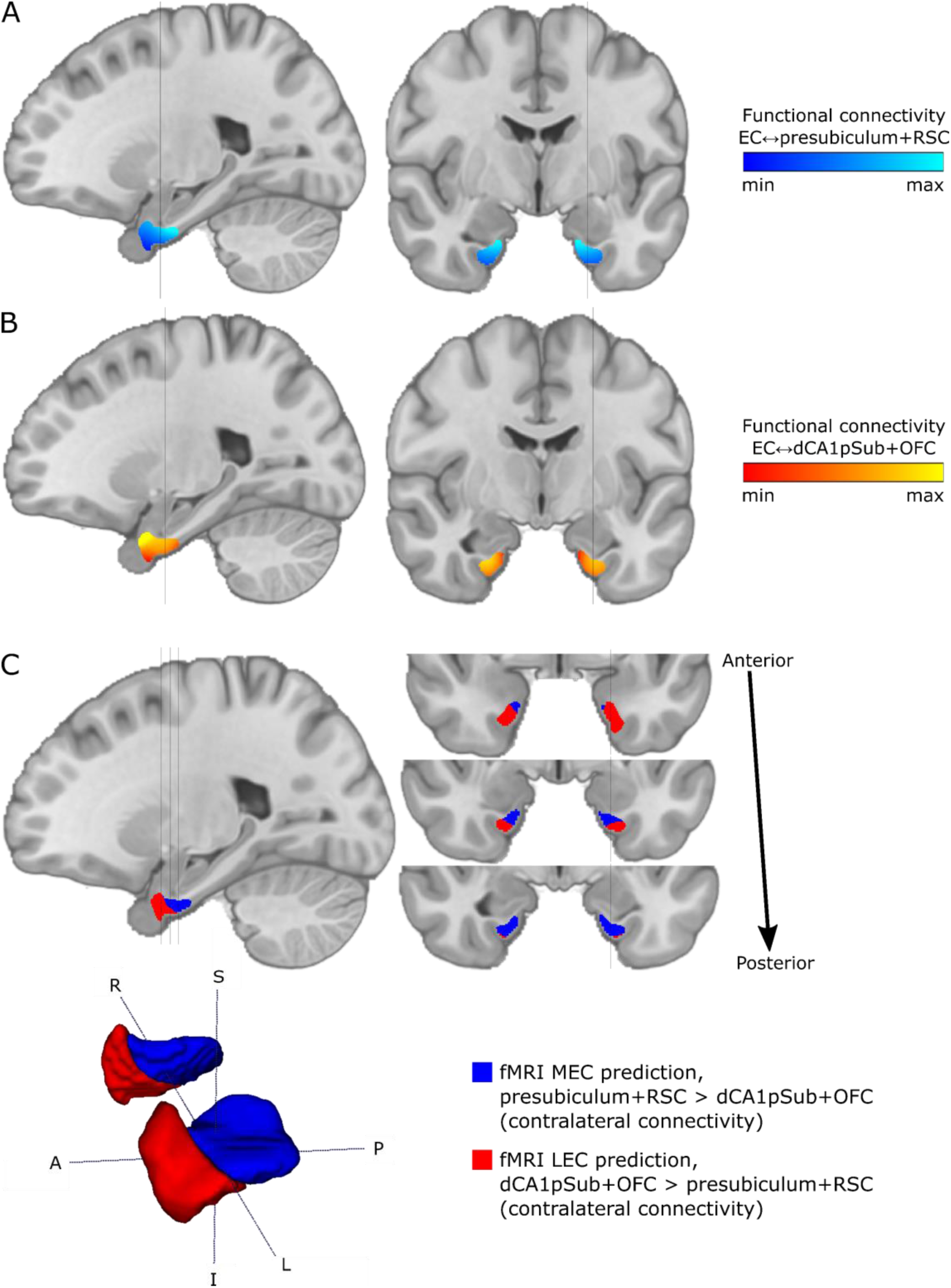
Group-averaged and phase encoding direction-averaged functional connectivity maps and MEC vs. LEC segmentations from contralateral rs-fMRI analysis. Connectivity maps in the right EC are seeded from ROIs in the left hemisphere, whereas the left EC is seeded from the right hemisphere. Results are shown on selected sagittal (left) and coronal (right) slices in MNI space. Grey lines on sagittal slices show level of adjacent coronal slices, and vice versa. **A:** EC connectivity with presubiculum+RSC, **B:** EC connectivity with dCA1pSub+OFC. **C:** Segmentation of MEC (blue) and LEC (red) homologues based on the connectivity maps, shown both on sagittal and coronal slices and in 3D (bottom row; both hemispheres are shown). The estimated degree of posterior-anterior orientation of the border for this segmentation was 68.0 ± 2.5 %, whereas the degree of ML orientation was 9.7 ± 2.3 %. S = superior, I = inferior, A = anterior, P = posterior, R = right, L = left.

**Supplementary Table 3:** Degree of posterior-anterior (PA) and medial-lateral (ML) orientation of the border between MEC and LEC for individual segmentations and resulting individual MEC/LEC size ratios. The degree of PA and ML orientation is given as a percentage between 0 and 100%, dependent on the angle between the MEC-LEC center of gravity vector and the pure PA or ML vector, respectively. For the individual segmentations, the sizes of the resulting MEC and LEC were not scaled to have equal sizes, and the MEC/LEC size ratios are therefore shown. All numbers are given as the mean ± standard deviation. As for the group-level segmentations, the individual segmentations show a larger degree of PA subdivision for fMRI than for DTI, and a larger degree of ML subdivision for DTI than for fMRI, although the % PA values are overall lower and the % ML values are higher than group-level results. This could be an effect of the unbalanced MEC/LEC size ratios where MEC tends to be larger than LEC, especially for the fMRI-based segmentations. 26 subjects were excluded from the fMRI comparisons due to the segmentation in the left and/or right hemisphere being classified as MEC only or LEC only, i.e. having an MEC/LEC size ratio of ∞ or 0.

| Individual segmentation approach |  | % Posterior-anterior (PA) axis | % Medial-lateral (ML) axis | MEC/LEC size ratio |
| --- | --- | --- | --- | --- |
| Presubiculum+RSC/dCA1pSub+OFC | DTI | 48.8 ± 13.0 | 32.0 ± 19.9 | 1.50 ± 0.69 |
| | fMRI | 57.0 ± 14.8 | 22.7 ± 19.8 | 5.17 ± 15.15 |

**Supplementary Figure 8:**
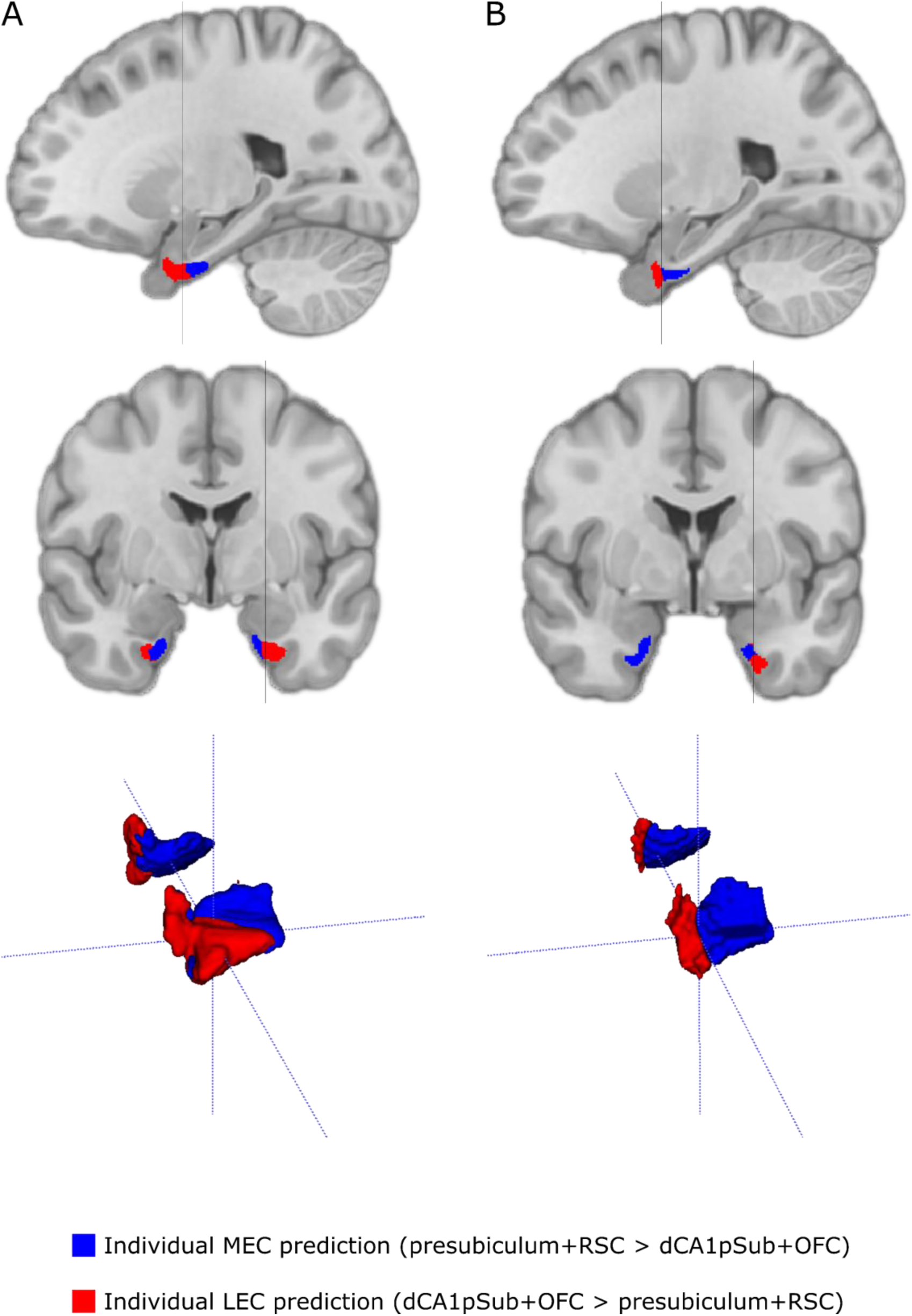
Individual DTI and fMRI segmentation results from an example subject. MEC is indicated by blue, and LEC is indicated by red. Upper row shows selected sagittal slices, middle row shows selected coronal slices, and bottom row shows the segmentations in 3D. Results are shown in MNI space. Grey line on sagittal slices shows level of coronal slice below, and vice versa. **A:** DTI-based segmentation results. The degree of posterior-anterior (PA) and medial-lateral (ML) subdivision for this segmentation is 55.6 % and 25.4 %, respectively. The MEC/LEC size ratio is 1.04. **B:** fMRI-based segmentation results. The degree of PA and ML subdivision for this segmentation is 49.5 % and 0.7 %, respectively. The MEC/LEC size ratio is 2.7.

**Supplementary Figure 9:**
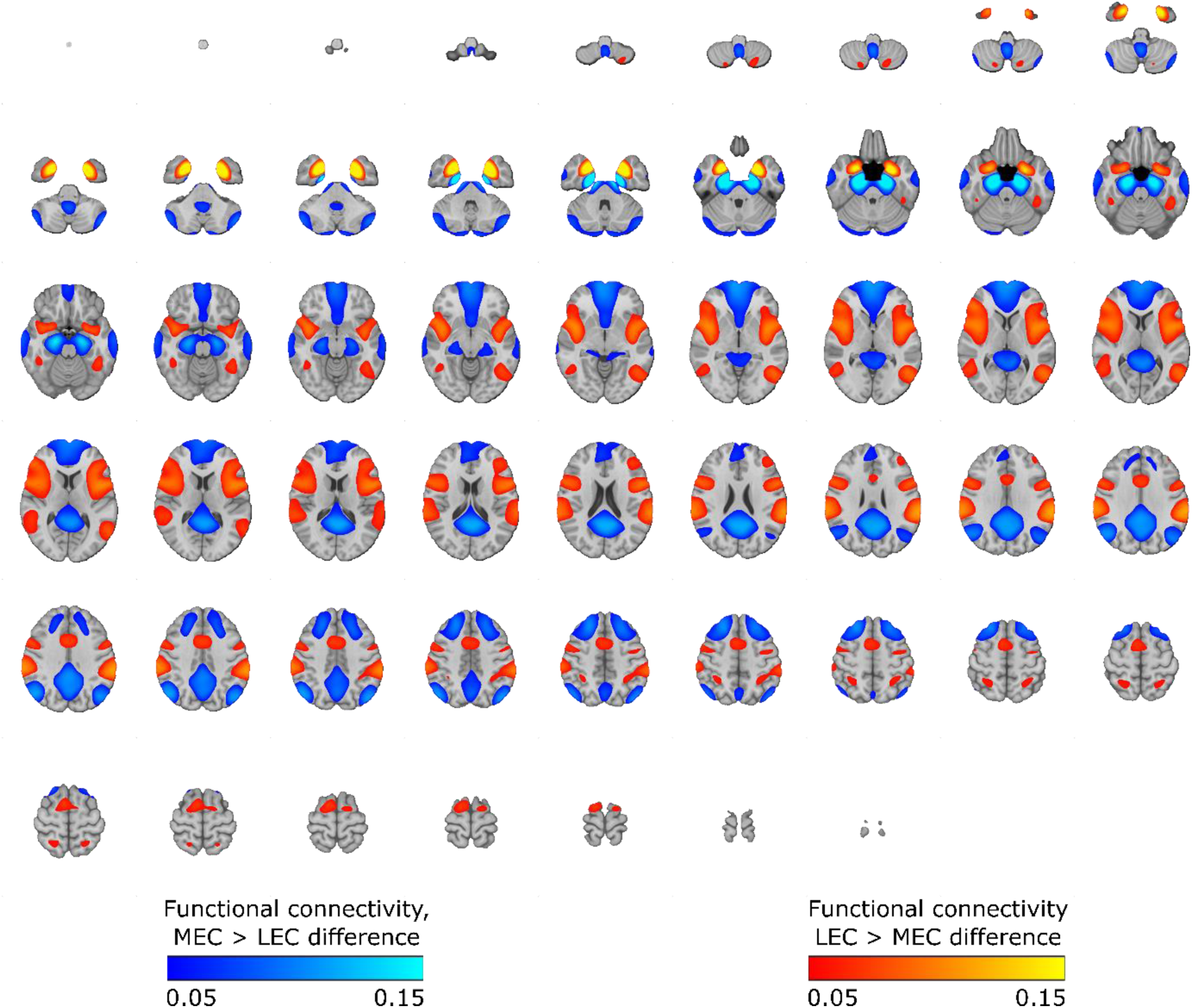
Group-averaged and phase encoding direction-averaged difference map of resting-state functional connectivity seeded from MEC and LEC ROIs from the DTI-based segmentation. Results are shown on axial slices throughout the brain. Blue means higher connectivity with MEC than with LEC, and red means higher connectivity with LEC than with MEC.

**Supplementary Figure 10:**
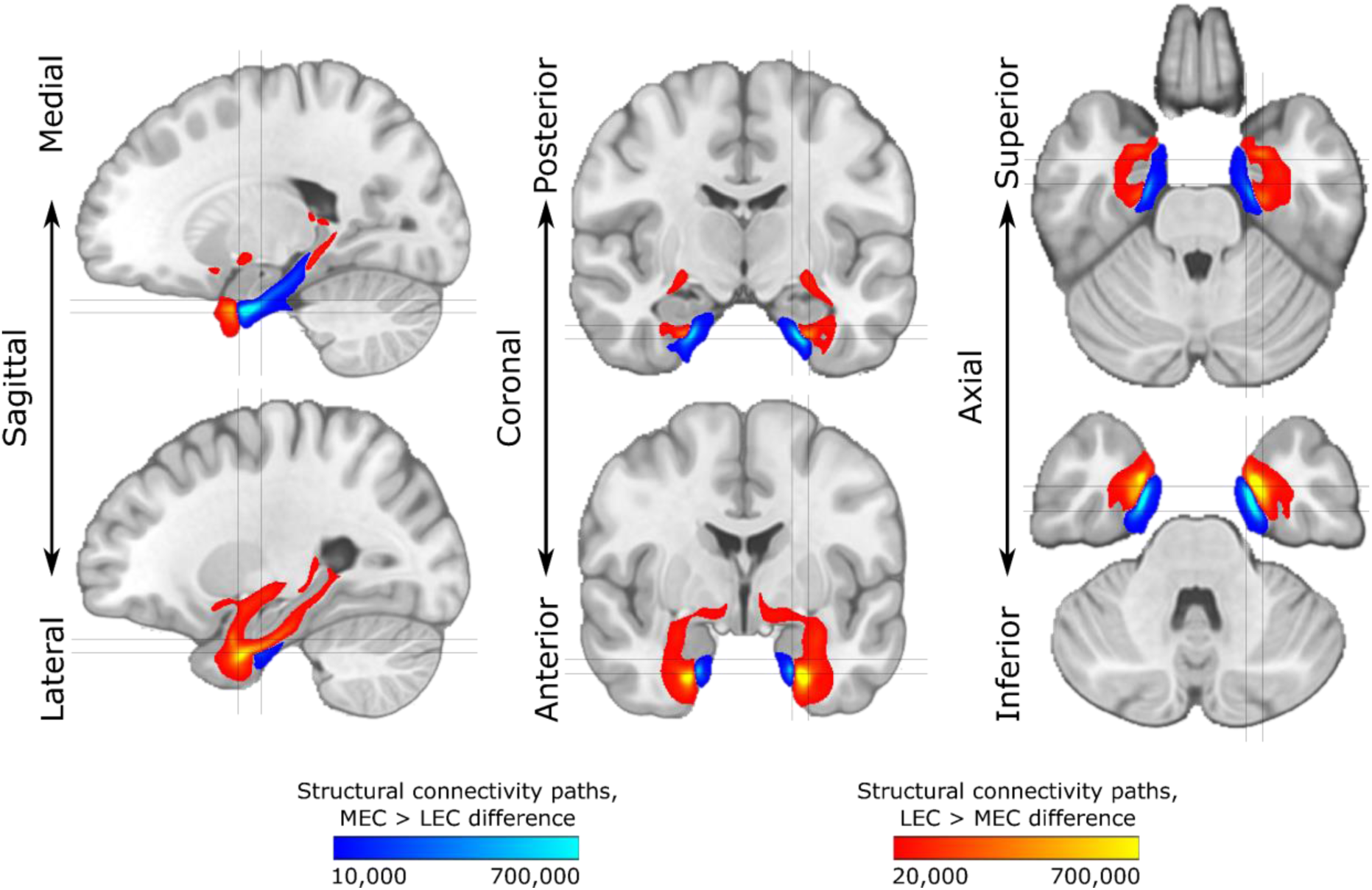
Group-averaged and field strength-averaged difference map of structural connectivity paths seeded from MEC and LEC ROIs from the fMRI-based segmentation. Results are shown on selected sagittal (left), coronal (middle) and axial (right) slices. Grey lines show levels of slices in the other planes. Blue means a higher probability that the path is connected with MEC than with LEC, and red means a higher probability that the path is connected with LEC than with MEC.

**Supplementary Table 4:** Summary of individual results from running DTI tractography seeded from MEC/LEC ROIs defined from group-level fMRI analysis, and from running fMRI connectivity analysis seeded from MEC/LEC ROIs defined from group-level DTI analysis. Values shown are mean ± standard deviation of results within presubiculum, dCA1pSub, RSC and OFC ROIs. For each ROI and for DTI and fMRI separately, connectivity measures from MEC vs. LEC seeds are compared. For “correctly cross-validated” results, i.e. that MEC > LEC connectivity for presubiculum and RSC ROIs or that LEC > MEC connectivity for dCA1pSub and OFC ROIs, the row is marked with green color. Statistically significant differences (p < 0.006; Bonferroni corrected for 8 multiple comparisons) are marked in bold.

|  | DTI |  | fMRI |  |
| --- | --- | --- | --- | --- |
|  | MEC | LEC | MEC | LEC |
| Presubiculum | 57199 ± 38709 | 56059 ± 35360 | **0.48 ± 0.11** | **0.35 ± 0.12** |
| dCA1pSub | **3394 ± 3329** | **8294 ± 6408** | **0.48 ± 0.11** | **0.40 ± 0.11** |
| RSC | 5695 ± 4382 | 4558 ± 3884 | **0.32 ± 0.11** | **0.24 ± 0.12** |
| OFC | **17 ± 42** | **83 ± 100** | **0.20 ± 0.10** | **0.22 ± 0.10** |

**Supplementary Figure 11:**
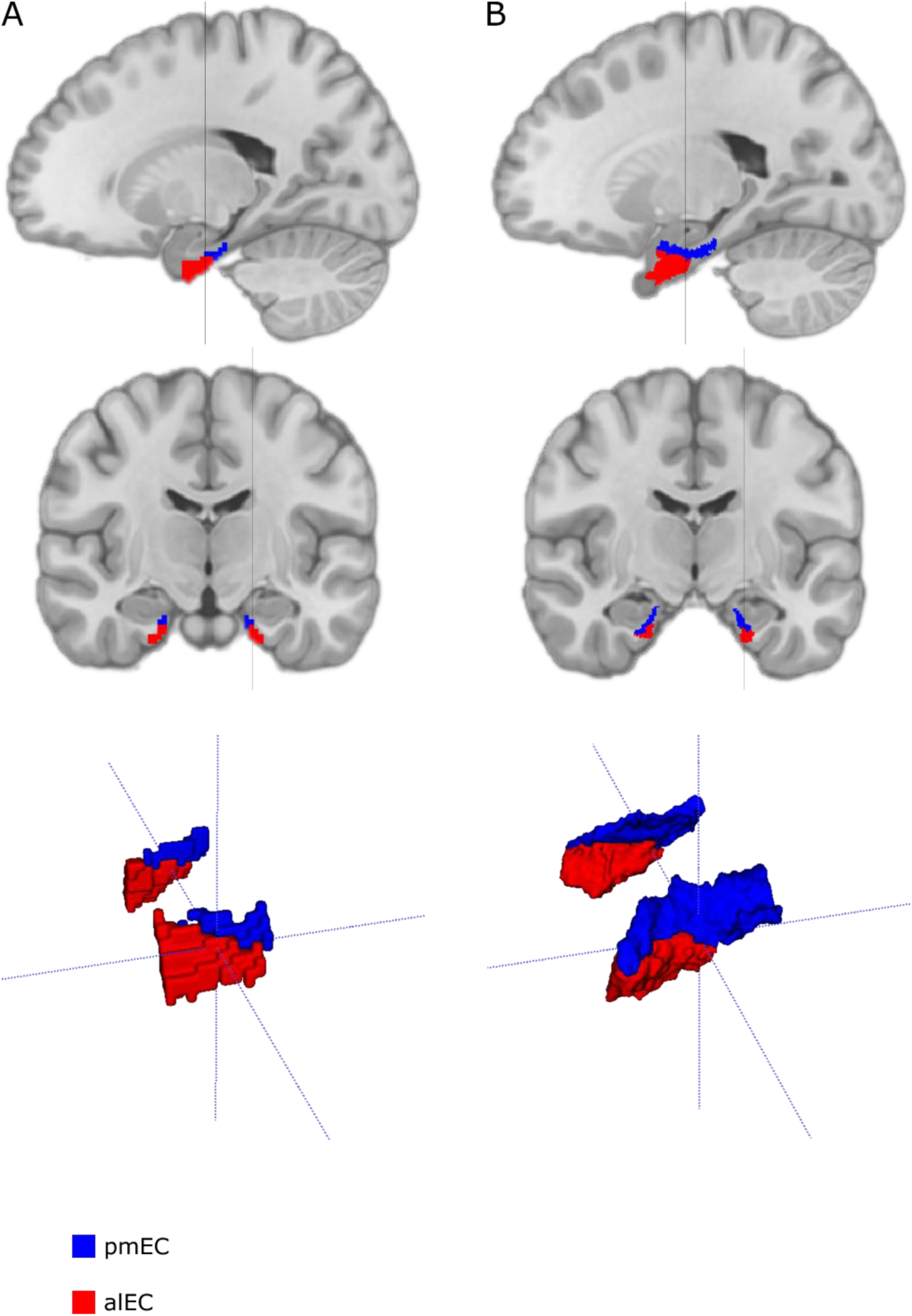
Posterior-medial EC (pmEC) and anterior-lateral EC (alEC) segmentation results from previous fMRI studies (A: Maass et al. eLife 2015, B: Navarro Schröder et al. eLife 2015). pmEC is indicated in blue, whereas alEC is indicated in red. Upper row shows selected sagittal slices, middle row shows selected coronal slices, and bottom row shows the segmentations in 3D. Results are shown in MNI space. Grey line on sagittal slices shows level of coronal slice below, and vice versa. As shown in Supplementary Table 2, these segmentations show a higher degree of posterior-anterior subdivision and a lower degree of medial-lateral subdivision than the segmentation results from the current study, although they look qualitatively similar to some extent. However, a direct comparison between the studies is difficult due to several factors: i) different seed regions have been used for the analyses; ii) different ROI delineations of the EC itself have probably been used; iii) the segmentations have different resolutions; and iv) there are uncertainties in the co-registration processes from study templates to MNI space, which would occlude direct anatomical comparison.

